# Virulence on Pm4 kinase-based resistance is determined by two divergent wheat powdery mildew effectors

**DOI:** 10.1101/2025.07.04.663163

**Authors:** Zoe Bernasconi, Aline G. Herger, Maria Del Pilar Caro, Lukas Kunz, Marion C. Müller, Ursin Stirnemann, Megan A. Outram, Victoria Widrig, Matthias Neidhart, Jonatan Isaksson, Seraina Schudel, Sebastian Rösli, Thomas Wicker, Kyle W. Bender, Cyril Zipfel, Peter N. Dodds, Melania Figueroa, Javier Sánchez-Martín, Beat Keller

## Abstract

The wheat resistance gene *Pm4* encodes a kinase fusion protein and has gained particular attention as it confers race-specific resistance against two major wheat pathogens: powdery mildew and blast. Here, we describe the identification of AvrPm4, the mildew avirulence effector recognised by Pm4, using UV-mutagenesis, and its functional validation in wheat protoplasts. We show that AvrPm4 directly interacts with and is phosphorylated by Pm4. Using genetic association and QTL mapping, we furthermore demonstrate that evasion of *Pm4* resistance by virulent mildew isolates relies on a second fungal component, *SvrPm4*, which suppresses *AvrPm4*-induced cell death. Surprisingly, *SvrPm4* was previously described as *AvrPm1a*. We show that SvrPm4, but not its inactive variant svrPm4, is recognised by the NLR immune receptor Pm1a. These multiple roles of a single effector provide a new perspective on fungal (a)virulence proteins and their combinatorial interactions with different types of immune receptors.

## Introduction

Wheat yields are severely impacted by pests and pathogens, accounting for over 20% of global losses (Savary *et al*., 2019). Resistance breeding is a key strategy for crop protection and reduction of losses. It often relies on dominant resistance (*R*) genes encoding immune receptors that recognise pathogen-delivered avirulence (Avr) effectors and activate immune responses, usually culminating in cell death (Dodds & Rathjen, 2010; Ngou *et al*., 2022). While most cloned *R* genes in wheat encode nucleotide-binding leucine-rich repeat (NLR) receptors, kinase fusion proteins (KFPs) are emerging as a distinct class of immune receptors specifically found in cereals (Chen *et al*., 2024). Much of our current understanding of KFPs comes from tandem kinase proteins—a major KFP subclass—which consist of two kinase domains, sometimes with additional domains of unknown function (Sánchez-Martín & Keller, 2021; Reveguk *et al*., 2025). Recent studies demonstrated that some tandem kinase proteins rely on a helper NLR to trigger effector-induced cell death, as shown for Sr62 in *Ae. tauschii* and RWT4 in wheat (Chen *et al*., 2025; Lu *et al*., 2025).

Beyond tandem kinase proteins, other KFPs, composed of at least one kinase domain and additional domains have been described in cereals, yet remain poorly understood (Fu *et al*., 2009; Liu *et al*., 2024). Among them, Pm4, a kinase-MCTP protein, is particularly notable: the *Pm4* gene located on wheat chromosome 2A encodes two alternative isoforms derived by alternative splicing, Pm4-V1 and Pm4- V2, both required for resistance to the biotrophic pathogen *Blumeria graminis* f. sp. *tritici* (*Bgt*) (Sánchez-Martín *et al*., 2021). *Pm4* occurs as several alleles (*Pm4a-g)* which encode highly similar proteins. *Pm4a* and *Pm4b* have been studied extensively in near-isogenic backgrounds, where they show partially overlapping race-specific resistance spectra against *Bgt* (Sánchez-Martín *et al*., 2021). Furthermore, recent work has shown that *Pm4* also confers resistance to the hemibiotrophic wheat blast pathogen *Magnaporthe oryzae* (Asuke *et al*., 2024; O’Hara *et al*., 2024). In fact, the wheat blast resistance gene *Rmg7* was found to be identical with *Pm4a*, whereas the *Rmg8* resistance gene on wheat chromosome 2B could be assigned to a *Pm4* homoeolog with identical sequence to *Pm4f* (Asuke *et al*., 2024). The corresponding wheat blast effector *AVR-Rmg8* was cloned earlier (Anh *et al*., 2018), and was recently shown to be recognised by multiple *Pm4* alleles, including *Pm4a*, *Pm4b* and *Pm4f* (Asuke *et al*., 2024; O’Hara *et al*., 2024). The KFP encoded by *Pm4* therefore represents a highly important resistance source in wheat with multiple alleles providing resistance to the obligate biotrophic *Bgt* and the hemibiotrophic wheat blast pathogen simultaneously.

Despite the progress made in describing novel KFPs, the mechanistic understanding of how such proteins induce immunity is yet to be established. Kinase activity in KFPs was suggested to be relevant for Avr effector recognition and/or downstream signalling, a fact further supported by the finding of loss-of-function mutants affected in the kinase domain of KFPs such as Pm4 (Sánchez-Martín *et al*., 2021). Moreover, the tandem kinase protein RWT4 (allelic to the powdery mildew R protein PM24;(Lu *et al*., 2020; Arora *et al*., 2023)) phosphorylates the avirulent but not the virulent variant of the wheat blast effector PWT4, suggesting that the phosphorylation of the Avr effector plays a key role in the resistance mechanism (Sung *et al*., 2025). For the barley stem rust resistance protein RPG1, two fungal proteins were shown to work synergistically to induce phosphorylation and trigger a hypersensitive response (HR) (Nirmala *et al*., 2011). These examples also highlight the importance of direct interaction between KFPs and their recognised effectors. Understanding these mechanisms will require the identification and characterisation of Avr effectors that interact with KFPs such as Pm4.

Methods based on genetic association, such as biparental mapping or genome-wide association studies, have been widely used to identify *Avr* loci in *Blumeria* and were combined with cell death assays in heterologous systems such as *Nicotiana benthamiana* and host protoplasts (Bourras *et al*., 2015; Saur *et al*., 2019; Kunz *et al*., 2023). More recently, AvrXpose, an approach based on UV-mutagenesis and selection of gain-of-virulence mutants, has also proven effective in identifying genes controlling avirulence in *Bgt* (Bernasconi et al., 2024). To date, all known *Avr* genes in *Bgt* encode small (100-150 amino acid residues), secreted effector proteins with a predicted RNase-like structure and have been shown to trigger a cell-death response upon recognition by corresponding wheat NLR immune receptors (reviewed in (Bilstein-Schloemer *et al*., 2025)). In contrast, no *Bgt* Avr effector recognised by a KFP has been identified so far.

Suppressors of *R*-gene-mediated immunity have been reported across bacterial, oomycete, and fungal pathogens (reviewed in (Wu & Derevnina, 2023)). Notable examples from phytopathogenic fungi include the *I* locus in *Melampsora lini* suppressing L7- and Lx-mediated resistance in flax (Dodds *et al*., 2004), *AvrLm4-7* in *Leptosphaeria maculans* suppressing *Rlm3* and *Rlm9*-mediated resistance in oilseed rape (Plissonneau *et al*., 2016; Ghanbarnia *et al*., 2018), and the *SvrPm3* effector in *Bgt* with the ability to suppress resistance mediated by the wheat NLR Pm3 (Bourras *et al*., 2015; Bourras *et al*., 2019). Most suppressors identified so far were found to suppress NLR-mediated immunity by diverse molecular mechanisms including interference with Avr recognition, NLR signalling or NLR homeostasis (Wu & Derevnina, 2023). In contrast, little is known about the existence or mode-of-action of suppressor proteins affecting non-NLR resistance proteins such as KFPs.

In this study, we describe the identification of *AvrPm4* from *Bgt* using UV mutagenesis and show that the encoded protein is recognised by the wheat Pm4 resistance protein, resulting in cell death. AvrPm4 is a non-canonical Avr effector that directly interacts with Pm4. Furthermore, using genetic association and QTL mapping, we describe the identification of *SvrPm4*, a suppressor of *AvrPm4*-induced cell death and show that SvrPm4 activity is the main mechanism by which *Bgt* evades *Pm4* resistance. Additionally, we show that SvrPm4, but not its inactive variant svrPm4, is recognised by the NLR Pm1a. Together, these findings represent a significant step forward in understanding the resistance mechanisms of *Pm4*, and open new avenues for studying multi-pathogen resistance provided by KFPs in cereals.

## Results

### Powdery mildew mutants that simultaneously gain virulence on *Pm4a* and *Pm4b* exhibit mutations in a novel type of effector gene

To identify the *Bgt* effector recognised by Pm4, we used the UV mutagenesis-based approach AvrXpose (Bernasconi *et al*., 2024) and mutagenised the reference *Bgt* isolate CHE_96224, avirulent on near-isogenic wheat lines containing *Pm4a* or *Pm4b*. Six mutants with gain of virulence on both *Pm4* alleles were identified (hereafter, *AvrPm4* mutants). By phenotyping the *AvrPm4* mutants on a set of resistant wheat tester lines, we observed that their gain of virulence was specific to *Pm4* and did not affect avirulence on other *Pm* genes (**Supp. Fig. S1**). Whole genome sequencing revealed that all six *AvrPm4* mutants had mutations in the coding sequence of the effector gene *Bgt-55142* (**Fig. 1A**). This was the only commonly mutated gene among all six *AvrPm4* mutants (**Supp. Tables S1-S2**), suggesting that *Bgt-55142* is *AvrPm4*. We therefore focused on *Bgt-55142* for further analyses.

**Fig. 1.**
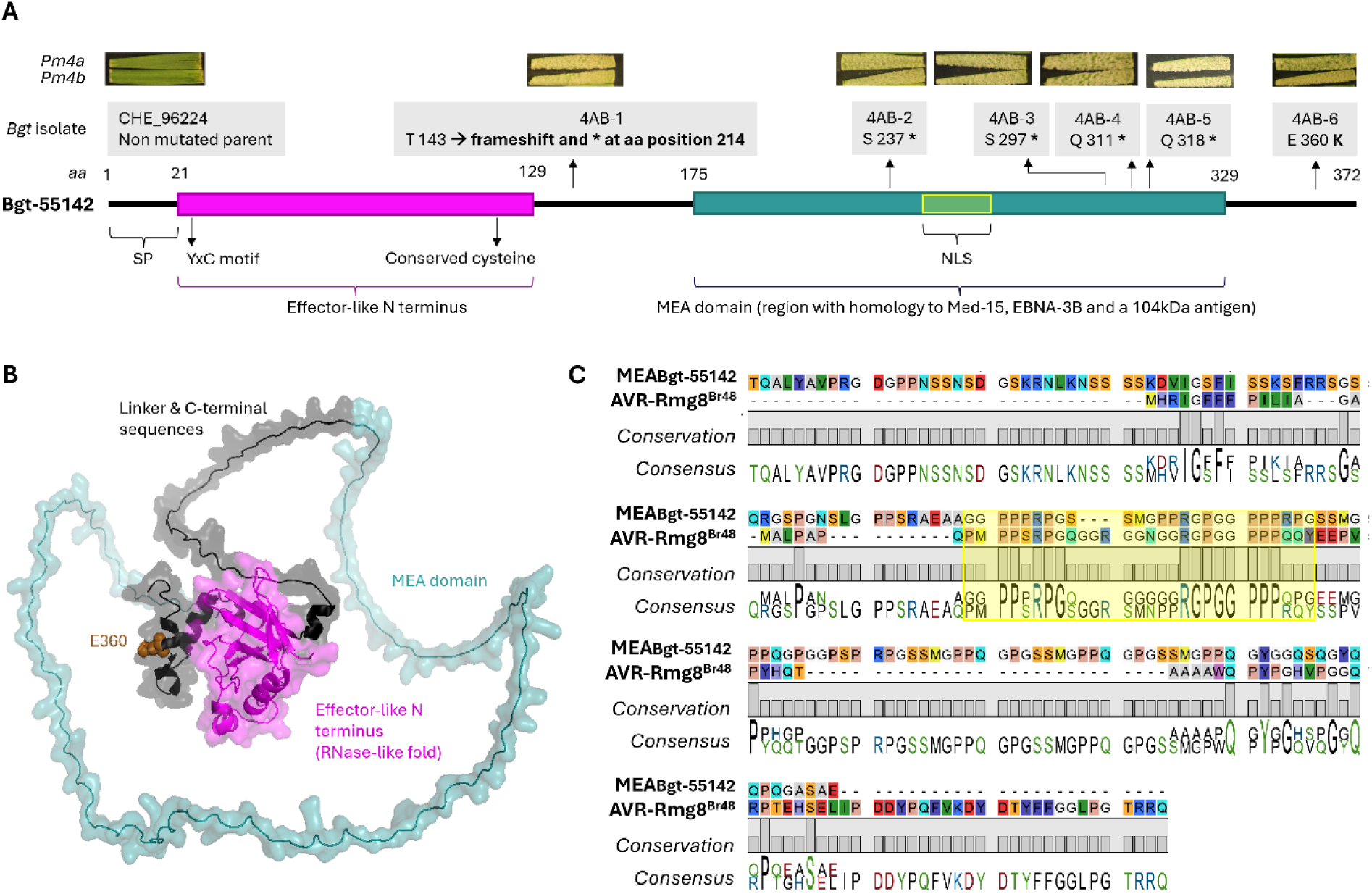
*Pm4* gain-of-virulence mutants all exhibit mutations in *Bgt-55142*, encoding a non-canonical effector. (**A**) Schematic representation of Bgt-55142 protein structure, indicating predicted domains (described in detail in **Supp. Fig. S2C**) and a nuclear localisation signal (NLS) predicted by NLStradamus. The phenotype and genotype of the *Bgt* gain-of-virulence mutants on *Pm4a* and *Pm4b* are shown above. (**B**) Cartoon and surface representation of the Alphafold3 structural prediction of Bgt-55142, with domains shown in different colours and labelled accordingly. Glutamate at position 360, mutated in mutant 4AB-6 to a lysine, is represented by spheres and indicated in brown. The predicted aligned error (PAE) is depicted in **Supp. Fig. S2A**. (**C**) Sequence alignment of the MEA domain of Bgt-55142 (amino acids 175 to 329) and AVR-Rmg8 reveals a region of high identity. In particular, the RGPGGPPP motif is 100% conserved, and overlaps with the NLS (in yellow; **Supp. Fig. S2B**).

Intriguingly, five *AvrPm4* mutants had nonsense mutations in *Bgt-55142* leading to a truncated protein or frameshifts, while only one (4AB-6) had a missense mutation leading to an amino acid change (E360K; **Fig. 1A-B**). *Bgt-55142* is highly expressed at the haustorial stage (Bourras *et al*., 2016; Praz *et al*., 2018) and encodes a protein with a signal peptide and an N-terminal domain with a predicted RNase-like structure, including the characteristic YxC motif found in all previously identified *Bgt* Avr proteins (**Fig. 1A**; (Bourras *et al*., 2016; Praz *et al*., 2018)). However, it differs from typical powdery mildew Avrs in length and structure, being considerably larger (372 amino acid residues) and containing a C-terminal domain, which is predicted with low confidence by Alphafold3, likely due to its repetitive and glycine and proline rich nature (**Fig. 1A-C, Supp. Fig. S2A-B**). Based on NCBI conserved domain search, this domain shows homology to a MED15 domain, an EBNA-3B domain, and a 104-kDa microneme/rhoptry antigen domain (**Supp. Fig. S2C**), the first two being previously shown to be implicated in transcriptional regulation (see also discussion; (Burgess *et al*., 2006; Malik & Roeder, 2010)). Given that homologies with these three domains overlap, we defined the C-terminus of Bgt- 55142 as a unique domain, which we renamed MEA (each letter representing one of the three identified hits; **Fig. 1A-B, Supp. Fig. S2C**). The MEA domain contains five repeated sequences (three complete, with 17 amino acids, and two incomplete ones, with 10 amino acids) as well as a predicted nuclear localisation signal (NLS) (**Supp. Fig. S2B-C**). Furthermore, the software DP-bind indicates a high probability of binding DNA, particularly in the MEA domain of Bgt-55142 (**Supp. Fig. S2D**).

The identity of the wheat blast effector AVR-Rmg8, recognised by Pm4 (Rmg8/Rmg7), is known (Anh *et al*., 2018; Asuke *et al*., 2024; O’Hara *et al*., 2024). We aligned its protein sequence with Bgt-55142 and found a region with high amino acid identity, corresponding to part of the MEA domain of Bgt- 55142, which also contains the predicted NLS. Notably, an eight-amino acid glycine-proline-rich motif (RGPGGPPP) is identical between the two effectors (**Fig. 1C**).

### *Bgt-55142^96224^* expression induces cell death in wheat protoplasts

To validate *Bgt-55142* as *AvrPm4*, we performed a cell death assay in wheat protoplasts, similar to the one performed for *AVR-Rmg8* (Asuke *et al*., 2024). Gene sequences of the avirulent and two mutated forms of *Bgt-55142* (*Bgt-55142^96224^*, *Bgt-55142^4AB-6^*, and *Bgt-55142^4AB-2^*), along with those of known control *Avrs* (*AVR-Rmg8*, *AvrPm3^a2/f2^*, and *AvrSr50),* were codon-optimised for expression *in planta* and cloned into expression vectors without signal peptide. Constructs were co-transfected with a YFP reporter into protoplasts of the transgenic ‘Bobwhite S26’ wheat line stably expressing *Pm4b-V1* and *Pmb4b-V2,* previously described (Sánchez-Martín *et al*., 2021). A non-transgenic sister line lacking *Pm4b* served as a negative control. We observed a strong reduction in relative YFP fluorescence in the *Pm4b*-overexpressing line upon co-expression with *Avr-Rmg8* and *Bgt-55142^96224^*, but not in the sister line (**Fig. 2A-B**). In contrast, neither of the two *Bgt-55142* mutant alleles *Bgt-55142^4AB-6^*and *Bgt- 55142^4AB-2^*, showed a reduction in relative YFP fluorescence nor did the negative controls *Avr-Pm3^a2/f2^* and *Avr-Sr50* (**Fig. 2A-B**). This indicates that *Bgt-55142^96224^*, from now on referred to as *AvrPm4^96224^*, is recognised in a *Pm4b*-dependent manner and subsequently triggers cell death. We then transfected *Pm4b-V1* and *Pm4b-V2* in protoplasts of the wheat line ‘Bobwhite S26’ (lacking *Pm4*), alone and in combination with each other. Interestingly, *Pm4b-V2* reduced fluorescence intensity by itself and in combination with *Pm4b-V1*, suggesting that it is partially autoactive (**Fig. 2C**). When transfecting both *Pm4b-V1* and *Pm4b-V2* in combination with *AvrPm4^96224^*, we observed a significant reduction in fluorescence intensity compared to *avrPm4^4AB-2^* (**Fig. 2C**), further confirming *Pm4b*-mediated cell death induction by *AvrPm4^96224^*in wheat. Co-expression of *AvrPm4^96224^* with *Pm4* in *N. benthamiana* did not lead to cell death, indicating that one or more additional genetic components, present in wheat but not in *N. benthamiana*, are needed for triggering *Pm4*-mediated immunity (**Supp. Fig. S3**).

**Fig. 2.**
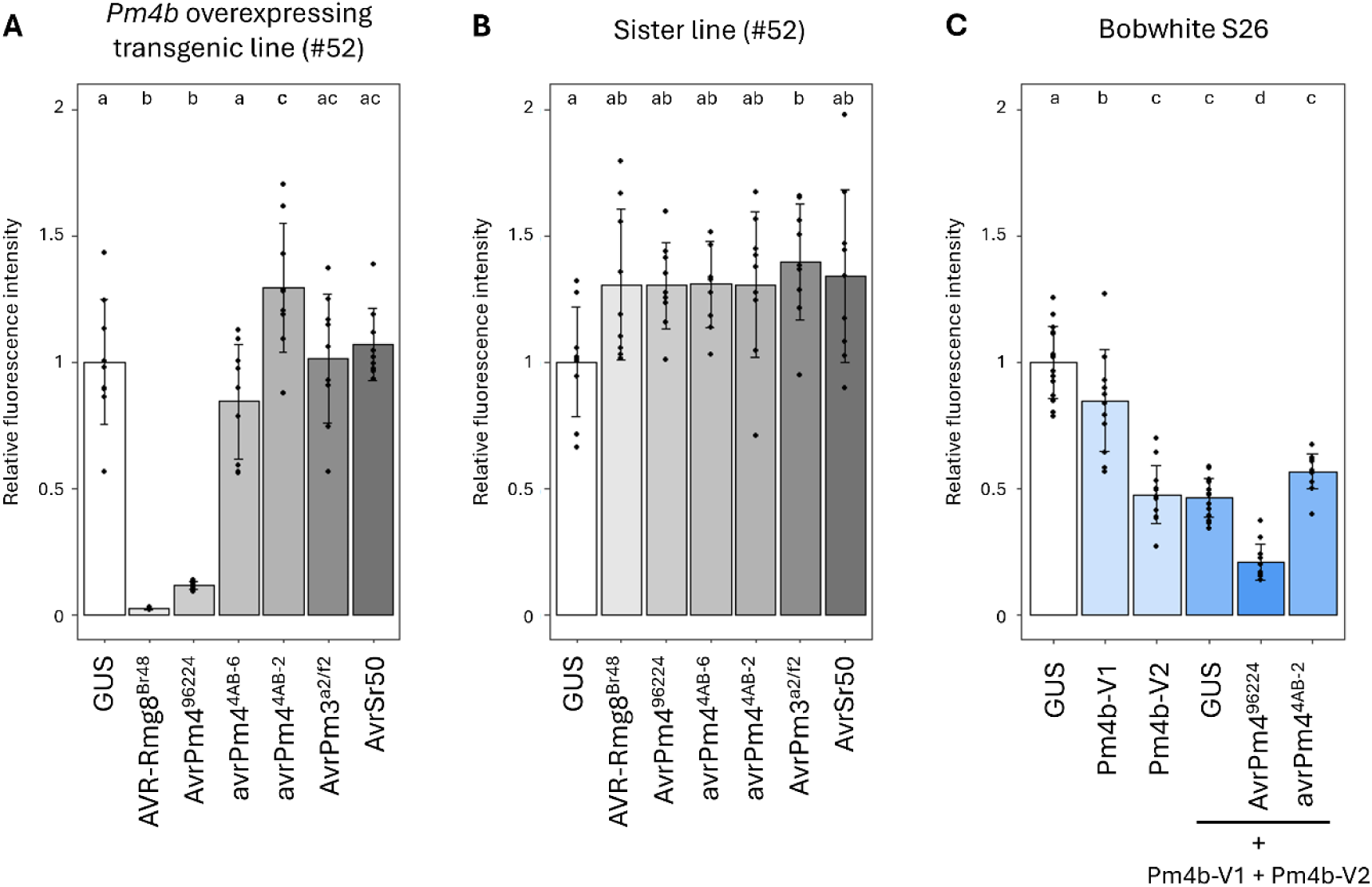
*Bgt-55142^96224^* induces cell death in *Pm4b*-containing wheat protoplasts. (**A**) Protoplasts generated from a transgenic line overexpressing *Pm4b* and (**B**) its *Pm4b*-free sister line were transfected with different *Avr* effectors. *Avr-Rmg8* was used as a positive control, while *AvrPm3^a2/f2^*and *AvrSr50* served as negative controls. In the presence of *Pm4b, Bgt-55142^96224^* (*AvrPm4^96224^*) induced cell death in wheat protoplasts in contrast to the mutant and truncated versions *Bgt-55142^4AB-2^* and *Bgt- 55142^4AB-6^*. (**C**) Transfection of *Pm4b-V1*, *Pm4b-V2* and *Bgt-55142^96224^* (*AvrPm4^96224^*), but not *Bgt-55142^4AB-2^* (*avrPm4^4AB-2^*), induced cell death in protoplasts of the wheat cultivar Bobwhite S26. *Pm4b-V2* shows autoactivity, leading to increased cell death in all tests containing *Pm4b-V2*. For all three panels, each treatment was assessed with two technical and three biological replicates, and the experiments were repeated three times (total n = 9 biological replicates per experiment). Statistical significance (*P* < 0.05) was determined using a two-sided analysis of variance (ANOVA) followed by a Tukey’s HSD test. Boxes with the same letter are not significantly different.

### Pm4 directly interacts with AvrPm4, resulting in its localisation to the ER network

Multiple tandem kinases were found to interact with their corresponding Avr effectors as part of their resistance function (Chen *et al*., 2025; Lu *et al*., 2025; Sung *et al*., 2025). We therefore investigated whether Pm4 binds AvrPm4 in a split luciferase assay in *N. benthamiana*. Interestingly, we found that both AvrPm4^96224^ and the truncated avrPm4^4AB-2^ interacted with both Pm4b isoforms, Pm4b-V1 and Pm4b-V2 (**Fig. 3A-B**). In contrast, we did not observe an interaction between Pm4 and the unrelated effector BgtE-5764 or between AvrPm4^96224^ and WTK4, another KFP, confirming the specificity of the AvrPm4/Pm4 interaction (**Fig. 3A-B**). We consistently observed a stronger interaction signal between AvrPm4 and Pm4b-V1 compared to Pm4b-V2, possibly reflecting differential protein accumulation between the isoforms (**Supp. Fig. S4A**). The fact that the virulent variant avrPm4^4AB-2^ also interacts with Pm4 suggests that the interaction alone is not sufficient to trigger an immune response.

**Fig. 3.**
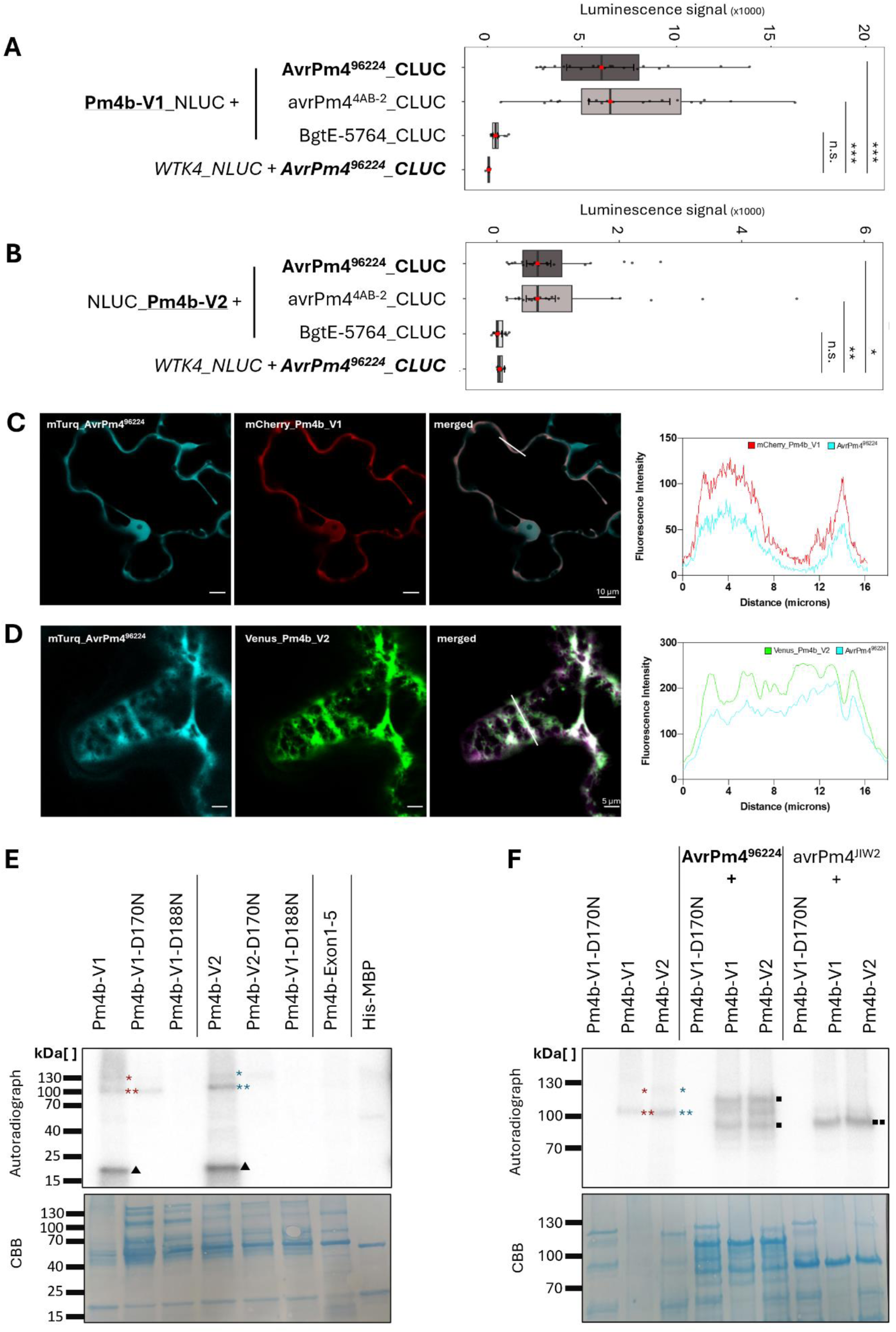
AvrPm4^96224^ interacts with both Pm4b isoforms V1 and V2. Split-luciferase assay showing the interaction of (**A**) NLUC tagged Pm4b-V1 and (**B**) NLUC tagged Pm4b-V2 with CLUC tagged AvrPm4^96224^ and CLUC tagged avrPm4^4AB-2^ in *N. benthamiana.* Co-expression of the two *Pm4b* isoforms with *BgtE-5764,* and of *AvrPm4^96224^* with *WTK4* served as negative controls. *AvrPm4^96224^* interacts with both *Pm4b* isoforms. Significance and *P* values (significance threshold of *P* < 0.05) were calculated with a two-sided analysis of variance (ANOVA) followed by multiple comparisons of means (contrast analyses) with Bonferroni correction (significance codes: *** <= 0.001, ** <= 0.01, * <= 0.05). (**C**) and (**D**) Subcellular co-localisation in *N. benthamiana* of mTurquoise-AvrPm4^96224^ and Pm4. When co-expressed with Pm4b-V1, AvrPm4^96224^ co-localises to the cytoplasm (C), Scale bar = 10 μm while co-expression with Pm4b-V2 redirects AvrPm4 to the ER. Scale bar = 5μm. (**E**) *In vitro* kinase assay showing autophosphorylation and transphosphorylation activity of purified Pm4 variants in the presence of [ɣ-32P]. All constructs shown have an MBP tag. All reactions were supplemented with the common substrate Myelin Basic Protein (MyBP, 21 kDa, indicated by a triangle). Constructs included *Pm4b-V1*, *Pm4b-V2ΔTMD*, and their kinase-dead mutants (D170N, D188N), as well as a truncated construct containing only the kinase domain (*Pm4b-Exon1–5*, 77.5 kDa). His-MBP alone served as negative control. Wild-type Pm4b-V1 (migrates at 106 kDa) and Pm4b-V2 (migrates at 107 kDa) showed kinase activity (indicated by a single *). Truncated degradation products migrating between 80-90 kDa (indicated by **) were also detected. Mutations D170N and D188N reduced or abolished kinase activity, respectively. Coomassie-stained PVDF membranes served as loading controls. (**F**) *In vitro* phosphorylation assay of AvrPm4^96224^ and avrPm4^4AB-2^ by Pm4. All constructs shown have an MBP tag. Pm4b-V1 and Pm4b-V2ΔTMD were incubated with AvrPm4^96224^ and avrPm4^4AB-2^ in the presence of [ɣ-32P]. AvrPm4^96224^ (110 kDa, with degradation products ranging between 70 kDa and 100 kDa (**Supp. Fig. S5B**), indicated by two single squares) and avrPm4^4AB-2^ (indicated by a double square) were phosphorylated by both Pm4b isoforms. As a control, the same AvrPm4 constructs were incubated with kinase dead Pm4b-V1-D188N.

As previously reported, Pm4b-V1, in the absence of Pm4b-V2, predominantly localises to the cytoplasm, whereas Pm4b-V2 or Pm4b-V1/Pm4b-V2 complexes localise to the endoplasmic reticulum (ER) (Sánchez-Martín *et al*., 2021). To explore whether AvrPm4 and Pm4b share similar localisation patterns indicative of a functional interaction, we assessed their subcellular localisation in *N. benthamiana*. We co-expressed mTurquoise-AvrPm4^96224^ with either mCherry-Pm4b-V1 or Venus- Pm4b-V2 and examined their subcellular localisation by confocal microscopy. Intensity plot analyses revealed that mTurquoise-AvrPm4^96224^ co-localises with mCherry-Pm4b-V1 in the cytoplasm (**Fig. 3C, Supp. Fig. S4B**), whereas co-expression with Venus-Pm4b-V2 re-localises AvrPm4^96224^ to the ER network (**Fig. 3D, Supp. Fig. S4B**). These findings provide additional evidence supporting the interaction between the two Pm4b isoforms and AvrPm4^96224^.

### Pm4 exhibits kinase activity and phosphorylates AvrPm4

Earlier work has shown that both Pm4 isoforms share a kinase domain that exhibits sequence characteristics of an active kinase. Furthermore, multiple loss-of-function mutants affecting this domain confirmed its essential role in resistance (Sánchez-Martín *et al*., 2021). Therefore, we tested whether Pm4 exhibits kinase activity and whether AvrPm4 is a substrate for phosphorylation by Pm4.

We established an *in vitro* assay by expressing full-length Pm4b-V1 (kinase domain + C2C domain) and a truncated, soluble version of Pm4b-V2 (Pm4b-V2ΔTMD, kinase domain + C2D domain, lacking the transmembrane and PRT-C domains) with an MBP (Maltose Binding Protein) tag in *E. coli*. Both isoforms were purified using amylose resin and protein integrity was assessed by Coomassie staining and western blotting (**Fig. 3E, Supp. Fig. 4C**). Pm4 showed some partial degradation (**Fig. 3E, Supp. Fig. 4C**).

Both Pm4b-V1 (106 kDa) and Pm4b-V2ΔTMD (107 kDa) exhibited *in vitro* kinase activity and autophosphorylation (**Fig. 3E**, single asterisk). A truncated fragment with a size of 80-90 kDa was also visible on autoradiographs (**Fig. 3E**, double asterisk), indicating that either the truncated form is catalytically active or a phosphorylation target of the full-length protein present in the extract. Importantly, both Pm4b isoforms also exhibited transphosphorylation activity, phosphorylating myelin basic protein (MyBP) as an exogenous substrate (**Fig. 3E**, triangle). A truncated construct containing only the kinase domain (exons 1-5, lacking a C2 domain) did not exhibit kinase activity, suggesting that the C2C domain in Pm4b-V1 and C2D domain in Pm4b-V2 are required for kinase activity (**Fig. 3E**).

We previously identified two EMS mutants compromised in *Pm4-*resistance to powdery mildew, each carrying a non-synonymous mutation in a conserved motif of the kinase domain: a D170N substitution in the HLDLKPAN motif of the catalytic loop, and a D188N substitution in the activation loop (Sánchez-Martín *et al*., 2021). To assess the functional impact of these substitutions, we introduced each substitution into both Pm4b isoforms and tested auto- and transphosphorylation activity using MyBP as a substrate. Auto- and transphosphorylation were strongly reduced by D170N and completely abolished by the D188N substitution (**Fig. 3E**), demonstrating a correlation between Pm4 kinase activity and resistance.

We further hypothesised that kinase activity may distinguish functional from non-functional *Pm4* allelic variants. To test this, we evaluated the *in vitro* kinase activity of the proteins encoded by alleles *Pm4a*, *Pm4f* and *Pm4g*. *Pm4a* has been described to have a largely overlapping resistance spectrum against *Bgt* with *Pm4b*, while *Pm4f* provides wheat blast resistance by recognising the wheat blast effector *AVR- Rmg8*. Importantly, *Pm4g* was reported to lack resistance activity against both pathogens and is thus considered a non-functional allele (Sánchez-Martín *et al*., 2021; Asuke *et al*., 2024; O’Hara *et al*., 2024). We observed that both Pm4a and Pm4f isoforms exhibited auto- and transphosphorylation activity comparable to Pm4b, whereas Pm4g did not show detectable kinase activity (**Supp. Fig. S5A**), further supporting the hypothesis that kinase activity is crucial for Pm4 resistance function.

Given the importance of kinase activity for *Pm4* resistance and cell death induction, along with its ability to transphosphorylate proteins such as MyBP, and its interaction and co-localisation with AvrPm4, we investigated whether Pm4 can directly phosphorylate AvrPm4. To this end, we expressed MBP-tagged AvrPm4^96224^ and the truncated virulent variant MBP-avrPm4^4AB-2^ in *E. coli,* followed by protein purification using amylose resin. We incubated the purified effectors with Pm4b-V1, Pm4b-V2ΔTMD and the kinase-inactive mutant Pm4b-V1-D188N as a negative control. Despite similar size ranges to Pm4b-V1 and Pm4b-V2, the MBP-AvrPm4^96224^ and MBP-avrPm4^4AB-2^ were clearly distinguishable from Pm4-derived bands (**Fig. 3F, Supp. Fig. S5B**). Importantly, we observed phosphorylation of AvrPm4^96224^ in the presence of Pm4b-V1 and Pm4b-V2ΔTMD but not in presence of the kinase dead mutant Pm4b-V1-D188N (**Fig. 3F**). Surprisingly, despite its inability to trigger cell death (**Fig. 2**), the virulent variant avrPm4^4AB-2^ was also phosphorylated by both Pm4 isoforms (**Fig. 3F**). These results confirm AvrPm4-Pm4 interaction *in vitro*, but also suggest that phosphorylation of the Avr alone is insufficient to trigger an immune response.

### *Bgt* virulence on *Pm4* is controlled by a suppressor locus on *Bgt* chromosome 8

Loss-of-function UV-mutants of *AvrPm4* result in virulence on both *Pm4a* and *Pm4b* (**Fig. 1A**). To elucidate the natural mechanism of *Bgt* race-specificity on *Pm4a* and *Pm4b* we phenotyped a set of 78 *Bgt* isolates selected from a worldwide collection (Sotiropoulos *et al*., 2022) on the near-isogenic lines (NILs) ‘W804/8*Fed’ (*Pm4b*) and ‘Khapli/8*CC’ (*Pm4a*) and their corresponding susceptible controls ‘Federation’ (Fed) and ‘Chancellor’ (CC). For most isolates, we observed identical virulence phenotypes on both *Pm4* alleles (**Supp. Table S3**), consistent with previous findings indicating that *Pm4b* and *Pm4a* have largely overlapping - although not identical - recognition spectra (Sánchez-Martín *et al*., 2021).

We next assessed the genetic diversity of *AvrPm4* in the same subset of 78 *Bgt* isolates. *AvrPm4* was present in all tested isolates, and it was largely conserved, with only 20 out of 78 isolates exhibiting single nucleotide polymorphisms (SNPs) in the coding sequence compared to the reference isolate CHE_96224 (**Supp. Table S3**). Importantly, no correlation was found between *AvrPm4* genotype and virulence phenotypes on *Pm4b* and *Pm4a* NILs. To rule out expression polymorphisms, we analysed available RNA-sequencing datasets and observed no significant variation in *AvrPm4* expression between *Bgt* isolates with contrasting phenotypes on *Pm4*, such as CHE_96224, CHE_94202 and ISR_7 (all avirulent on *Pm4b/Pm4a*) versus GBR_JIW2 (virulent on *Pm4b*/*Pm4a*) (**Supp. Fig. S6, S7A**) (Praz *et al*., 2018; Kunz *et al*., 2023). Taken together, these observations suggest that additional genetic components beyond *AvrPm4* control virulence on *Pm4*. To identify such additional components, we used two independent approaches, namely genome-wide association study (GWAS) and bi-parental QTL mapping.

The above-mentioned *Bgt* collection of 78 isolates, phenotyped on the NILs ‘W804/8*Fed’ (*Pm4b*) and ‘Khapli/8*CC’ (*Pm4a*), provided the basis for GWAS. Using an avirulent isolate as a reference (CHE_96224), GWAS revealed two significant genetic associations for virulence on *Pm4b*, a strong association at the end of chromosome 8 (Chr-08), spanning from 10,704,663 to 10,864,069 bp in the reference assembly of CHE_96224, and a less pronounced association at the beginning of Chr-04 (2,736 to 457,765 bp) (**Fig. 4A**). GWAS analysis on *Pm4a* revealed a single significant genetic association located on Chr-08 (10,721,247 to 10,841,388 bp), overlapping with the locus identified for *Pm4b* (**Fig. 4B**).

**Fig. 4.**
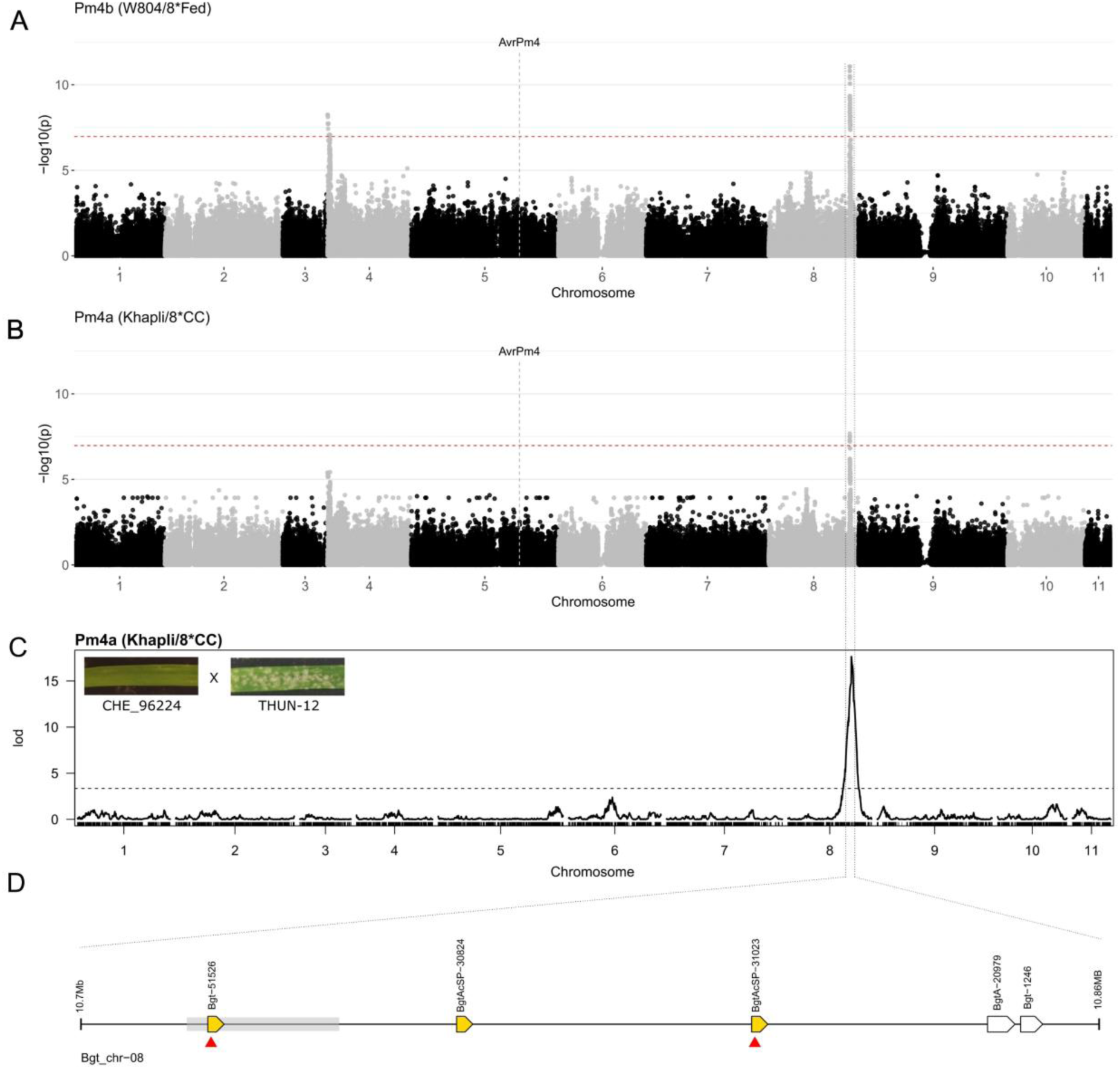
GWAS and QTL mapping identify a common locus on *Bgt* Chr-08 controlling virulence on *Pm4b* and *Pm4a*. (**A, B**) GWAS analysis using 473,887 SNPs across the 11 *Bgt* chromosomes, based on phenotypes of 78 isolates on the *Pm4b* NIL ‘W804/8*Fed (**A**) and the *Pm4a* NIL ‘Khapli/8*CC’ (**B**). The red line indicates the significance threshold at *p* < 0.05 after Bonferroni correction. The genomic position of *AvrPm4* is indicated by a dashed line. (**C**) Result of a QTL analysis on *Pm4a* NIL ‘Khapli/8*CC, using 118 F1 progenies of the bi-parental mapping population CHE_96224 (avirulent) X THUN-12 (partially virulent). The virulence phenotypes of the parental isolates on the *Pm4a* NIL are shown in the top left corner. The dashed line indicates the significance threshold LOD (logarithm of the odds) value at *p* < 0.05, determined by 1,000 permutations. (**D**) Depiction of the genomic locus on Chr-08 identified by GWAS and QTL analysis. Locus borders were defined based on the GWAS analysis on the *Pm4b* NIL (W804/8*Fed). Candidate secreted effector genes are represented by yellow arrows, and non-effector genes by white arrows. Red triangles indicate candidate effectors exhibiting non-synonymous sequence polymorphisms between *Pm4b/Pm4a* virulent and avirulent reference isolates. The grey box indicates the location of the 13 best associated SNPs with the phenotype on *Pm4b* (W804/8*Fed) with an identical p-value of 8.660015e^-12^. Gene names are based on the gene annotation published for *Bgt* reference isolate CHE_96224 (Müller *et al*., 2019). Gene models are not drawn to scale.

To complement the GWAS approach, we performed QTL mapping on *Pm4a,* using a bi-parental cross (originally described in (Müller *et al*., 2019)) between the *Pm4a/b*-avirulent *Bgt* isolate CHE_96224 and the *B. g. triticale* isolate THUN-12, which displays partial virulence on *Pm4a* but is avirulent on *Pm4b* (**Fig. 4C**). To do so, we phenotyped 118 F_1_ progenies on the *Pm4a* NIL ‘Khapli/8*CC’ and conducted a QTL analysis using 119,023 genetic markers between the parental isolates. Remarkably, a single QTL was identified on Chr-08, spanning from positions 9,931,004 to 10,932,716 in the CHE_96224 reference assembly, overlapping with the Chr-08 locus detected in the GWAS analysis (**Fig. 4C**).

Consistent with our previous observation that genetic diversity within *AvrPm4* does not correlate with virulence phenotypes on *Pm4b* or *Pm4a*, neither GWAS nor QTL mapping identified a significant genetic association with the *AvrPm4* gene located on Chr-05 (**Fig. 4A-C**). Instead, both complementary approaches provide strong evidence for the presence of a major genetic component on Chr-08 controlling *Bgt* virulence on *Pm4b* and *Pm4a* resistance alleles. Given that loss-of-function mutations in *AvrPm4* alone are sufficient to result in virulence on *Pm4b* and *Pm4a* (**Fig. 1A**), it is unlikely that the locus on Chr-08 contains an additional *Avr* component. We therefore hypothesised that this region contains a suppressor of avirulence (*SvrPm4*), which interferes with AvrPm4 recognition by Pm4.

### *SvrPm4* (*Bgt-51526^JIW2^*) suppresses *AvrPm4/Pm4*-induced cell death in wheat protoplasts

We defined *SvrPm4* candidate genes within the identified locus on Chr-08 of the *Pm4*-avirulent isolate CHE_96224 based on the GWAS analysis on *Pm4b* (**Fig. 4D**). Focusing on effector genes with a predicted signal peptide, we identified three candidate genes within this interval: *Bgt-51526*, *BgtAcSP- 30824* and *BgtAcSP-31023*. Analysis of their expression levels during infection in four RNA-sequenced *Bgt* reference isolates exhibiting differential phenotypes on *Pm4b* and *Pm4a* (avirulent: CHE_96224, CHE_94202, ISR_7; virulent: GBR_JIW2) revealed no major expression polymorphisms (**Supp. Fig. 7A-B**). We therefore focused on sequence polymorphisms between avirulent and virulent reference isolates to define promising *SvrPm4* candidates. Whereas *BgtAcSP-30824* did not exhibit any sequence polymorphisms, both *Bgt-51526* and *BgtAcSP-31023* were consistently polymorphic between the *Pm4*- virulent reference isolate GBR_JIW2 and the avirulent isolates CHE_96224, CHE_94202 and ISR_7. We found that the effector protein Bgt-51526^JIW2^ differs by 27 amino acids from Bgt-51526^96224^, whereas BgtAcSP-31023^JIW2^ exhibits two amino acid polymorphisms and a premature stop codon at position 142 of the effector protein, compared to BgtAcSP-31023^96224^, found in avirulent isolates (**Fig. 5A**). We therefore considered both genes as *SvrPm4* candidates and proceeded to test their ability to suppress *Pm4*-mediated cell death in wheat protoplasts.

**Fig. 5.**
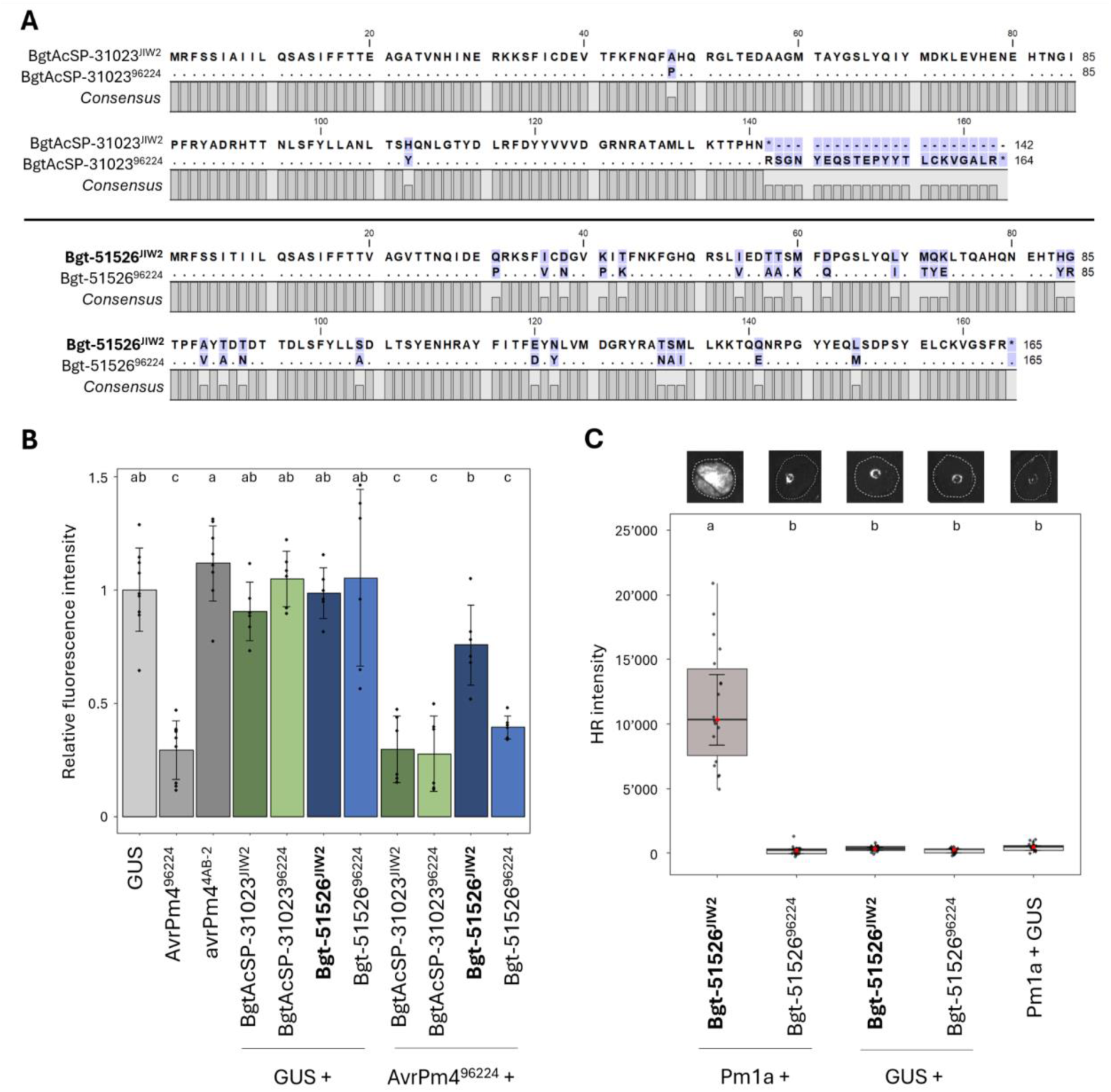
*AvrPm4^96224^*induced cell death in *Pm4*-containing wheat protoplasts is suppressed by co-expression of *SvrPm4^JIW2^*. (**A**) Protein sequence alignment of SvrPm4 candidates BgtAcSP-31023 and Bgt-51526 haplovariants found in the *Pm4* virulent isolate GBR_JIW2 and in the *Pm4* avirulent isolate CHE_96224. GBR_JIW2 is shown as the reference, with polymorphic residues in CHE_96224 highlighted. (**B**) Wheat protoplast assay showing suppression of *AvrPm4*-induced cell death by *SvrPm4^JIW2^*. Protoplasts isolated from the transgenic wheat line #52, overexpressing *Pm4b* (used in **Fig. 2**) were transfected with each *Svr* candidate individually or co-transfected with *AvrPm4*. Presumed suppressor-active candidates (*BgtAcSP-31023^JIW2^* and *Bgt-51526^JIW2^*) and suppressor-inactive candidates (*BgtAcSP-31023^96224^* and *Bgt-51526^96224^*) were tested. The experiment was performed twice with three biological replicates each (n = 6 total). (**C**) HR response in *N. benthamiana* upon *Agrobacterium*-mediated co-expression of *Bgt-51526^JIW2^* or *Bgt-51526^96224^* and *Pm1a-HA*, or a negative GUS control. The assay was performed three times with n = 6 leaves per experiment (n = 18 total). For all experiments, significance and *P* values (significance threshold of *P* < 0.05) were calculated with a two-sided analysis of variance (ANOVA) followed by a Tukey’s HSD test (**Supp. Table S3**). Boxes with the same letter are not significantly different.

We codon-optimised the two candidate genes without signal peptide and co-transfected them with a YFP reporter into the transgenic wheat line #52 stably expressing *Pm4b-V1* and *Pmb4b-V2,* as described above. In the absence of *AvrPm4*, none of the *SvrPm4* candidates triggered cell death in the *Pm4* transgenic line, consistent with our initial hypothesis that the locus on Chr-08 does not contain an *Avr* but rather an *Svr* factor (**Fig. 5B**). However, co-expression of *Bgt-51526^JIW2^* with *AvrPm4* significantly reduced the cell death response, whereas *Bgt-51526^96224^* or any of the two *BgtAcSP-31023* variants had no detectable suppressive effect (**Fig. 5B**). We therefore concluded that *Bgt-51526^JIW2^* (hereafter *SvrPm4^JIW2^*) indeed is a suppressor of *Pm4*-mediated cell death whereas *Bgt-51526^96224^*(hereafter *svrPm4^96224^*) is unable to suppress *Pm4* activity.

Among 78 tested *Bgt* isolates, we found extensive sequence variation within the *SvrPm4* gene, identifying 10 highly polymorphic haplovariants (**Supp. Fig. S8A, Supp. Table S3**). Importantly, all isolates carrying the inactive variant svrPm4^96224^ exhibit an avirulent phenotype on *Pm4b* and *Pm4a*, whereas all isolates carrying SvrPm4^JIW2^ are virulent on both *Pm4* alleles (**Supp. Fig. S8B**), highlighting the importance of the SvrPm4 activity in overcoming *Pm4* resistance. We identified multiple other haplovariants that were exclusively present in either avirulent or virulent isolates, thereby likely representing additional inactive and active SvrPm4 variants, respectively (**Supp. Fig. S8B**). Intriguingly, we also found three haplovariants associated with *Bgt* isolates showing differential phenotypes on *Pm4b* and *Pm4a*, suggesting that *SvrPm4* allelic variation may contribute to the subtle differences in *Pm4b* and *Pm4a* recognition spectra (**Supp. Fig. S8B**).Taken together, our data suggest a genetic two-component system controlling virulence on *Pm4* in natural *Bgt* isolates, in which the ability to overcome *Pm4*-mediated resistance is independent of the *AvrPm4* genotype but rather controlled by *SvrPm4* activity.

### The active suppressor SvrPm4^JIW2^ but not svrPm4^96224^ is recognised by the wheat NLR Pm1a

The gene *Bgt-51526* (*SvrPm4*) encodes an RNase-like effector protein that was previously identified as an Avr recognised by the NLR immune receptor encoded by the wheat resistance gene *Pm1a* (Kloppe *et al*., 2023). To determine whether *SvrPm4^JIW2^* (defined as Bgt-51526_2 variant by (Kloppe *et al*., 2023)) and the previously undescribed *svrPm4^96224^* variant induce differential HR responses, we co-expressed each variant with *Pm1a* in *N. benthamiana.* Consistent with previous findings, we observed a strong HR response upon co-expression of *SvrPm4^JIW2^* and *Pm1a,* confirming this variant’s avirulence toward *Pm1a*. Interestingly, co-expression of *svrPm4^96224^* and *Pm1a* did not result in an HR response (**Fig. 5C**). Thus, we conclude that *SvrPm4^JIW2^* is simultaneously an active suppressor of *Pm4* and an *Avr* of *Pm1a*, whereas the inactive suppressor *svrPm4^96224^* escapes *Pm1a* recognition. This observation suggests that combining *Pm4* and *Pm1a* in the same wheat genotype could provide complementary and potentially more durable resistance towards *Bgt* due to mutually excluding activities.

## Discussion

Avirulence on *Pm4* is controlled by at least two genes in *Bgt*. *AvrPm4* is an avirulence factor and its disruption results in virulence on both *Pm4a* and *Pm4b*. Furthermore, *AvrPm4*-triggered immunity is suppressed by *SvrPm4*. This multi-component system was uncovered using UV-mutagenesis, GWAS and QTL mapping, illustrating how the combination of gene identification tools allows the dissection of genetically complex systems in biotrophic fungal plant pathogens such as *Bgt*.

Previously identified Avr effectors in *Bgt* encode small (100-155 amino acid residues) RNase-like effectors recognised by NLR receptors (Bilstein-Schloemer *et al*., 2025). In contrast, AvrPm4, recognised by the KFP Pm4, is a 372-residue chimeric protein with an N-terminal RNase-like domain and a C-terminal MEA domain and differs substantially from previously identified *Bgt* Avr proteins.

Recent studies revealed that the *Pm4* resistance gene corresponds to previously identified wheat blast resistances *Rmg7* and *Rmg8*, and established that Pm4a, Pm4b and Pm4f can recognise the wheat blast effector AVR-Rmg8 (Asuke *et al*., 2024; O’Hara *et al*., 2024). Interestingly, AVR-Rmg8 (110 residues) is considerably smaller than AvrPm4 and lacks an RNase-like domain (Anh *et al*., 2018). However, it shares a nearly identical sequence motif with the MEA domain of AvrPm4 (**Fig. 1C**) suggesting the importance of this domain for Avr recognition by Pm4/Rmg8. Interestingly, we found that the MEA domain of AvrPm4 shows homologies to MED15, a crucial coactivator in eukaryotic transcription, and to EBNA-3B from the Epstein-Barr virus, which is also implicated in transcriptional control (**Fig. 1, Supp. Fig. S2,** (Krauer *et al*., 1998; Burgess *et al*., 2006; Malik & Roeder, 2010)). Additionally, the AvrPm4 MEA domain has a predicted nuclear localisation signal and a high probability of binding DNA (**Supp. Fig. S2C**). Thus, it is likely that this domain functions in transcriptional regulation in the host and contributes to *Bgt* virulence. Indeed, both *AvrPm4* and *AVR-Rmg8* are conserved within the gene pool of *Bgt* and wheat blast respectively, thereby indicating an important role of these effectors for pathogen fitness. Considering that the MEA domain might also be implicated in Avr recognition, as suggested by the nearly identical sequence motifs shared by AvrPm4 and AVR-Rmg8, this might represent a significant trade-off between preservation of effector function and evasion of Pm4 recognition for the fungal pathogen.

Mutation or loss of Avr effectors is a common mechanism to evade *R*-gene mediated immunity in fungal plant pathogens (Petit-Houdenot & Fudal, 2017). Loss of Avr effectors with crucial virulence functions might, however, result in reduced pathogen fitness. Therefore, multiple fungal plant pathogens were found to secrete suppressor proteins to mask Avr effectors or suppress NLR-mediated immunity thereby allowing to retain important Avr effectors (Wu & Derevnina, 2023). Previous studies in *Bgt* identified SvrPm3, an RNase-like effector that suppresses recognition of AvrPm3 effectors by NLR-type Pm3 resistance proteins (Bourras *et al*., 2015; Bourras *et al*., 2019). Like SvrPm3, the SvrPm4 effector belongs to the group of RNase-like effectors. Our study reveals that a member of this large effector group suppresses recognition by a KFP immune receptor. This underscores the diversity of molecular activities of these effectors and their importance as suppressors of race-specific resistance.

Intriguingly, a study in the blast pathogen (*Magnaporthe* spp.) has identified the *PWT4* effector, naturally present in oat-infecting blast isolates, as a suppressor of *Pm4* (*Rmg8*) when expressed in wheat blast (Inoue *et al*., 2021). This finding further highlights the similarities in genetic control of avirulence/virulence on *Pm4* between the powdery mildew and blast pathosystems.

Our GWAS and QTL analyses indicate that *SvrPm4* is the main factor controlling virulence on *Pm4a* and *Pm4b* within the global *Bgt* population (**Fig. 4**, **Supp. Fig. S8**). The importance of the *SvrPm4* locus is further corroborated by the fact that a genetic association between this locus and Pm4 resistance in wheat was also described in a recent study using host-pathogen biGWAS (Xie *et al*., 2025). Importantly, besides the active SvrPm4^JIW2^ and the inactive svrPm4^96224^ variants, we found a staggering array of additional SvrPm4 variants within the *Bgt* population that so far remain untested for their ability to suppress *Pm4*. Phenotype-genotype correlations suggest that additional *SvrPm4* variants may act as active suppressors against *Pm4* (**Supp. Fig. S8**), while others might target only a subset of *Pm4* alleles, thereby likely explaining the subtle differences in recognition spectra provided by different *Pm4* alleles ((Sánchez-Martín *et al*., 2021), this study). Moreover, considering this vast diversity of the *SvrPm4* gene it will be also important to establish whether suppression of *Pm4* and evasion of *Pm1a* recognition are indeed mutually exclusive, which would allow the efficient generation of disruptive selection pressures on this effector through combination of *Pm4* and *Pm1a* in wheat breeding programs.

The GWAS analysis on *Pm4b* revealed a second minor locus residing on chromosome 4. However, the same locus did not show a significant association with virulence on *Pm4a* and was also not identified in our QTL mapping analysis, indicating that it represents a minor or *Pm4b*-specific factor. Future studies should dissect the complex interactions between the largely conserved *AvrPm4*, the large diversity of *SvrPm4* and their consequences on *Pm4* suppression as well as the potential existence of additional minor factors.

The *Pm4* gene exhibits alternative splicing resulting in two isoforms Pm4-V1 and Pm4-V2 sharing an identical kinase domain, both necessary for resistance. Pm4b-V1 localises to the cytoplasm but to re- localises to the ER membrane upon co-expression with Pm4b-V2 (Sánchez-Martín *et al*., 2021). We found that AvrPm4 interacts with both isoforms and re-localises from the cytoplasm to the ER membrane upon co-expression with Pm4b-V2 (**Fig. 3A-D**). Furthermore, we found evidence for auto- and transphosphorylation activities of both Pm4 isoforms that result in AvrPm4 phosphorylation (**Fig. 3E-F**). The importance of Pm4 kinase activity for resistance is furthermore highlighted by the observation that all *Pm4* alleles with documented resistance activity against either *Bgt* or wheat blast (namely *Pm4a*, *Pm4b*, *Pm4f*) exhibit kinase activity while the non-functional *Pm4g* allele does not. Strikingly, when investigating the ability of Pm4 to phosphorylate AvrPm4, we also observed phosphorylation of the non-recognised, truncated avrPm4^4AB-2^ variant. This suggests that AvrPm4 phosphorylation alone is not decisive for the initiation of *Pm4*-mediated cell death. This finding contrasts with a recent report of Sung et al. (2025) that observed phosphorylation of the Avr protein PWT4 in presence of the KFP RWT4, but did not detect phosphorylation of a non-recognised PWT4 variant (Sung *et al*., 2025). These differences could indicate that molecular resistance mechanisms among KFPs are diverse, similar to the functional diversity observed for NLR-type immune receptors.

The wheat KFPs *Sr62* and *WTK3* were recently found to depend on a NLR protein for HR induction (Chen *et al*., 2025; Lu *et al*., 2025). Consistent with the existence of another wheat component, *Pm4* and *AvrPm4* co-expression in the heterologous *N. benthamiana* system failed to trigger HR (**Supp. Fig. S3**). We therefore hypothesise that *Pm4*, like other KFPs, relies on another host protein, likely an executor NLR conserved in the wheat gene pool, for cell death induction. Identifying this unknown component should be a priority of future studies to mechanistically understand the induction of *Pm4*-mediated resistance. Additionally, the striking parallels between *Bgt* and wheat blast pathogens and their Avr and Svr effectors in the context of *Pm4/Rmg8* resistance provides the unique opportunity to study the complex molecular events during *Pm4*-mediated immunity in two evolutionary distant pathosystems.

With a quickly increasing number of identified and characterised R/Avr pairs, both in powdery mildew and wheat blast, a complex molecular network also involving suppressors, modifiers and additional resistance genes start to be unravelled. In fact, besides *Pm4/Rmg8/Rmg7*, recent studies showed further parallels between cereal powdery mildew and wheat blast pathosystems. The wheat *RWT4* resistance gene, identified as a key host specificity factor against the blast pathogen, is allelic to the *Bgt* resistance gene *Pm24* (Arora et al., 2023) and the barley powdery mildew resistance gene *MLA3* was found to recognise the wheat blast effector *PWL2* (Brabham et al., 2024). Moreover, our findings that the active suppressor of Pm4, *SvrPm4^JIW2^,* is recognised by the NLR *Pm1a*, while the inactive *svrPm4^96224^* evades recognition, further highlights the complexity of the R-Avr molecular networks even within the wheat powdery mildew pathosystem. In conclusion, our study advocates for a shift from single-gene strategies to resistance gene stacking, combining kinase-based receptors like *Pm4* with NLRs such as *Pm1a*. Additionally, we propose joint wheat breeding programs for mildew and blast resistance to efficiently use overlaps and synergies between resistance sources resulting in multilayered and potentially more durable pathogen resistance.

## Materials & Methods

### Fungal, plant material and phenotyping experiments

The *Bgt* reference isolate CHE_96224 and the mapping population CHE_96224xTHUN-12 have previously been described in (Müller *et al*., 2019). The 78 *Bgt* isolates used for virulence phenotyping and GWAS analysis were selected from the worldwide diversity panel described in (Sotiropoulos *et al*., 2022). Isolates were maintained clonally on leaf segments of the susceptible wheat cultivar ‘Kanzler’ (K-57220) placed on food grade agar (0.5% PanReac AppliChem, Darmstadt, Germany) supplemented with 4.23 mM benzimidazole (Merck, Darmstadt, Germany; CAS 51-17-2) (Parlange *et al*., 2011).

The NILs Khapli/8*CC/8*Fed (*Pm4a*), Khapli/8*CC (*Pm4a*) and W804/8*Fed (*Pm4b*) were previously described (Sánchez-Martín *et al*., 2021). The *Pm4b*- overexpressing-transgenic line (event #52), first reported by Sánchez-Martín et al. (2021), was self-fertilised to the T4 generation, producing T5 homozygous Pm4b transgenic plants. A segregating, transgene-free sisterline (S#52) was used as a control. For virulence specificity testing of *Bgt* mutants, NILs with specific resistance genes were used: Axminster/8*Chancellor (*Pm1a*; (Hewitt *et al*., 2020)), Federation*4/Ulka (*Pm2a*; (Sánchez-Martín *et al*., 2016)), Asosan/8*Chancellor, Chul/8*Chancellor, Sonora/8*Chancellor, Kolibri and Michigan Amber/8*Chancellor (*Pm3a, Pm3b, Pm3c, Pm3d* and *Pm3f*, respectively; (Brunner *et al*., 2010)), transgenic Pm3e#2 (*Pm3e*; (Koller *et al*., 2019)), Kavkaz/4*Federation (*Pm8*; (Hurni *et al*., 2013)), USDA_Pm24 (*Pm24*), and “Amigo” (1AL.1RS translocation line, *Pm17*; (Singh *et al*., 2018)). Additionally, *Ae. tauschii* accessions TOWWC087, TOWWC112 and TOWWC154 were included (*WTK4*; (Arora *et al*., 2019)).

*Bgt* virulence phenotyping was performed using the primary leaf of 8-12 days old wheat seedlings, grown at 16 h light/8 h dark cycles, 18°C and 60% humidity. Leaves were cut into 3 cm fragments and placed on Petri dishes with 0.5% water-agar containing 4.23 mM benzimidazole (Merck, Darmstadt, Germany; CAS 51-17-2). Spores were collected with a funnel and blow-inoculated onto the Petri dish. The phenotype was then evaluated by eye as percentage of leaf coverage by mycelium (0-100%) after 6-8 days.

### Mutagenesis and mutant analyses

*Bgt* mutants were isolated as described previously (Bernasconi *et al*., 2024). Briefly, spores of the avirulent isolate CHE_96224 were irradiated with UV light and propagated on cv. Kanzler three times, before using them for infecting *Pm4a* and *Pm4b* containing NILs (see above). Colonies growing on *Pm4a* or *Pm4b* NILs were isolated from single spores and their virulence phenotype was confirmed through an infection test. Finally, spores were collected, frozen in liquid nitrogen, ground using a tissue homogeniser and DNA was extracted using a modified CTAB method (as described previously; (Bourras *et al*., 2015)). Library preparation and Illumina sequencing were performed at the Functional Genomics Center Zurich (Zurich, Switzerland) and Novogene (Cambridge, UK) using NovaSeq 6000 technology, as described previously (Bernasconi *et al*., 2024).

The raw reads obtained were trimmed using Trimmomatic v. 0.39 (Bolger *et al*., 2014), then mapped to the reference genome of the *Bgt* isolate CHE_96224, version 3.16 (Müller *et al*., 2019) using bwa mem v0.7 (Li & Durbin, 2009). The last steps involved sorting, removing duplicates and indexing the bam files, using Samtools v1.9 (Li *et al*., 2009). Haplotype calling, transposable element insertions and gene deletions or duplications were performed as described by (Bernasconi *et al*., 2024). As performed in our previous study, variants less than 1.5 or 2 kb away (for SNPs and TE insertions, respectively) from annotated genes were considered for further investigations (**Supp. Tables S1-S2**).

### Confocal imaging

Live-cell imaging was performed using a Stellaris 5 inverted confocal laser scanning microscope (Leica Microsystems, Wetzlar, Germany) equipped with an external fluorescence light source (Lumencor LED3, Beaverton, OR, USA), a 405 nm diode laser, and a Leica white light laser (WLL). Confocal imaging was conducted as previously described by (Sánchez-Martín *et al*., 2021), with minor modifications. Briefly, four-week-old *N. benthamiana* plants were infiltrated with *Agrobacterium tumefaciens* carrying the plasmids of interest. Three days post-infiltration, 5 × 5 mm leaf samples were mounted between a glass slide and a coverslip in a drop of water. Fluorescence was observed using the following excitation and emission settings: mTurquoise: Excitation at 405 nm; emission collected between 425–522 nm; Venus: Excitation at 515 nm; emission collected between 520–550 nm; mCherry: Excitation at 587 nm; emission collected between 592–652 nm. Fluorescence intensities in the cytosol and endoplasmic reticulum (ER) were measured using the Plot Line plugin in Fiji software (https://fiji.sc/). All experiments were conducted under strictly identical confocal acquisition parameters, including laser power, gain, zoom factor, resolution, and emission detection settings, ensuring minimal background noise and avoiding pixel saturation. Pseudo-coloured images were generated using the “Green,” “Magenta,” and “Turquoise” look-up tables (LUT) in Fiji.

### Cloning of expression constructs

The coding sequences of *AvrPm4^96224^, avrPm4^4AB-2^, avrPm4^4AB-6^, SvrPm4^JIW2^*, *SvrPm4^96224^* and *BgtE- 5764_96224_* lacking the signal peptide (as detected by SignalP4.0, (Petersen *et al*., 2011)), were codon-optimised for wheat expression using the codon-optimisation tool from Integrated DNA Technologies (IDT; Coralville, IA, USA; https://eu.idtdna.com) and subsequently gene-synthesised by our commercial partner BioCat GmbH (Heidelberg, Germany; https://www.biocat.com). Codon-optimised *SvrPm4* variants were cloned into the Gateway-compatible pENTR plasmid using In-Fusion cloning (Takara Bio, Kusatsu, Japan), then mobilised into the binary expression vector pIPKb004 (Himmelbach *et al*., 2007) using LR Clonase II (Invitrogen, Carlsbad, CA, USA). The pIPKb004-Pm1a-HA construct has been previously described (Hewitt *et al*., 2020). *AvrPm4^96224^, avrPm4^4AB-2^, BgtE-5764_96224_* and *WTK4* were cloned into the Gateway-compatible pDONR207 plasmid and then into the destination vectors for the split luciferase assay (GW_NLUC, NLUC_GW, GW_CLUC, CLUC_GW; (Gehl *et al*., 2011)) using LR Clonase II (Invitrogen, Carlsbad, CA, USA), while LUC tagged Pm4b-V1 and Pm4b- V2 have been previously described (Sánchez-Martín *et al*., 2021). For wheat protoplast expression all previously described constructs were cloned into the pTA22 vector (Arndell *et al*., 2024). Additionally, *avrPm4^4AB-6^, AvrSr50* (Chen *et al*., 2017), *AvrPm3^a2/f2^* (Bourras *et al*., 2015), *Pm4b-V1*, *Pm4b-V1- D188N*, *Pm4b-V2*, *Pm4b-V2-D188N*, as well as *GUS* were PCR amplified, cloned into the pDONR207 vector and subsequently transferred into the binary expression vector pTA22 using LR Clonase II (Invitrogen, Carlsbad, CA, USA). *YFP-pTA22* was used as described in (Arndell *et al*., 2024).

The coding sequences for *mTurquiose-AvrPm4^96224^* was codon-optimised for plant expression using the optimisation tools provided by Integrated DNA Technologies (IDT; Coralville, IA, USA; https://eu.idtdna.com). Synthesis of mTurquoise*-AvrPm4^96224^*, *mCherry_Pm4b-V1* and *Venus_Pm4b-V2* as well as cloning into the Gateway-compatible vector pDONR221 were performed by Life Technologies Europe BV (Thermo Fisher Scientific, Bleiswijk, Netherlands). These constructs were subsequently transferred into the binary expression vector pIPKb004 (Himmelbach *et al*., 2007) using LR Clonase II enzyme mix (Invitrogen, Carlsbad, CA, USA).

The CDS of *Pm4b-V1*, *Pm4b-V1-D170N*, *Pm4b-V1-D188N*, *Pm4a-V1, Pm4f-V1, Pm4g-V1* (full-length variants), as well as *Pm4b-V2-dTMD*, *Pm4b-V2-D170N-dTMD*, *Pm4b-V2-D188N-dTMD, Pm4a-V2, Pm4f-V2, Pm4g-V2* (kinase + C2D domain), *AvrPm4^96224^* and *avrPm4^AB22^* were PCR amplified and cloned in frame into the vector pMAL-C4E between the restriction sites *EcoRI* and *BamHI,* in order to generate fusion proteins carrying a N-terminal maltose binding protein (MBP) tag (Walker *et al*., 2010). Ligation of the PCR product and vector backbone was performed using T4 DNA Ligase (New England Biolabs, Ipswich, MA, USA). The His-MBP Exon 1-5 coding sequence was PCR amplified using primers PC_90 and PC_91 with flanking *BsaI* recognition sites and unique overhangs to enable directional and seamless assembly. The construct was assembled using the Golden Gate cloning system, using type IIS restriction enzymes and T4 DNA ligase in a one-tube reaction. The assembled product was cloned into the destination vector pET28a(+)-GG.

The constructs were verified by full plasmid sequencing (Microsynth AG, Balgach, Switzerland). Primers used for cloning are listed in **Supp. Table S4**. All codon-optimised effector and resistance gene sequences used in this study are listed in **Supp. Table S5.** All expression constructs were transformed into the *Agrobacterium tumefaciens* strain GV3101 by electroporation.

### Cell death assay in wheat protoplasts

For the cell death assay in wheat protoplasts, we followed the description in (Arndell *et al*., 2024). Briefly, *Avr-*genes and *R-*genes were cloned into the plasmid pTA22 without tag and subsequently extracted using the Xtra Midi Plus Endotoxin-free MidiPrep Kit Kit (Macherey-Nagel, Düren, Germany). The DNA was eluted in ultra-pure water. Transgenic seedlings of susceptible cultivar Bobwhite S26 overexpressing Pm4b (#52) and its respective sister line (S#52) were grown in a chamber under a cycle of 12 h light (100 µmol m^−2^ s^−1^) and 12 h dark at 24 °C for 7 days. The protoplasts were isolated by peeling off the epidermis and digesting the leaf in a cellulase RS and macerozyme R-10 (Onozuka; Yakult Honsha, Tokyo, Japan) solution. After the purification steps described in (Arndell *et al*., 2024) the protoplast solution was diluted to a concentration of 3 x 10^5^ cells per mL. In a 2 mL tube the different plasmids were mixed for transfection. Three picomoles (3 pmol) of the plasmids containing *YFP* and the individual *Avrs* were used for transfection. Due to Pm4b-V2 autoactivity and to maintain an equal ratio between Pm4b-V1 and Pm4b-V2, 0.5 pmol of plasmids containing *Pm4b-V1, Pm4b-V1- D188N, Pm4b-V2* and *Pm4b-V2-D188N* were used. Where one DNA component is missing, the amount of DNA was compensated with a GUS expressing construct. 200 μL of protoplasts were then added to the DNA together with a PEG solution (volume PEG solution = 200 μL + volume of DNA mix = app. 230 μL; 40% w/v PEG-4000, 0.2 M mannitol, 100 mM CaCl_2_). After a gentle homogenisation, the transfection reaction was stopped by adding W5 solution (2 mM MES-KOH, 5 mM KCl, 125 mM CaCl_2_, 154 mM NaCl). All the reagents used for preparing the solutions were from Sigma-Aldrich (St. Louis, MO, USA). After transferring the protoplasts into a 12-well cell culture plate, they were incubated at 23°C for 16-20 hours in the dark. The next day, YFP fluorescence was measured (Excitation: 500 nm, Emission 541 nm) with two technical replicates and three biological replicates per treatment using a microplate reader (Synergy H1, BioTek Instruments, Winooski, VT, USA). Each experiment was repeated two to three times as indicated in the figure legends. All the measurements were normalised to the average of values from the GUS transfected protoplasts for each experiment individually.

### *Agrobacterium*-mediated gene expression in *N. benthamiana*

*Agrobacterium*-mediated expression in *N. benthamiana* was performed as described by (Bourras *et al*., 2019). In brief, *Agrobacterium tumefaciens* (strain GV3101) were grown in liquid Luria Broth (LB) medium containing appropriate antibiotics overnight at 28°C. For *N. benthamiana* infiltration, cultures were briefly washed in LB medium and resuspended in infiltration medium (200 µM Acetosyringone, 10 mM MgCl2, 10 mM MES-KOH pH5.6) to an OD_600_ of 1.2 for HR testing, and to an OD_600_ of 1 for split-luciferase and co-localisation experiments, subsequently incubated at 28°C for two to four hours and finally infiltrated into leaves of 3-4 week old *N. benthamiana* plants.

### Split-luciferase assay

All the constructs co-infiltrated were previously mixed in a 1:1:1 ratio with the third component being p19-silencing-suppressor strain (Jay *et al*., 2023). Transient expression by agroinfiltration in *N. benthamiana* was performed as described above. At 3 dpi, 6-mm leaf discs were collected from each leaf and incubated in buffer containing 10 mM MES-KOH (pH 5.6) and 10 mM MgCl₂. Subsequently, 1 mM luciferin (BioVision, Milpitas, CA, USA) in 0.5% DMSO was added, and after 5 minutes incubation, luminescence was measured for 25 minutes (200 ms per well) using the LUMI imaging system (Tecan, Männedorf, Switzerland). The sum of the values per well was then used for statistical analyses. Two technical replicates and at least sixteen biological replicates were measured for each treatment, for a total of three experiments.

### Recombinant protein expression, purification and *in vitro* kinase assays

The N-terminally MBP tagged constructs in the *pMAL-c4E* vector (Walker *et al*., 2010) were transformed into BL21 pLYsS chemically competent Rosetta cells. 10 mL of an overnight culture was added to 1 L of LB media, the respective antibiotic and 20% glucose to an OD of 0.6 to 0.8. Subsequently protein expression was induced with 300 mM IPTG and grown overnight shaking at a reduced temperature of 18 °C.

After harvesting the cells by centrifugation at 5000 RPM for 20 minutes the pellet was resuspended in 40 mL purification buffer (50 mM HEPES-KOH (pH 7.2), 5% glycerol and 1x protease inhibitor tablets (cOmplete EDTA-free; Roche, Basel, Switzerland) and lysed by sonication (4 x 20 seconds with 40 seconds breaks at high intensity). Next the lysed cells were centrifuged at 35,000 g for 30 min at 4 °C. The supernatant was further incubated with washed amylose resin (New England Biolabs, Ipswich, MA, USA) and an addition of 2 mM DTT and 300 mM NaCl. After 30 minutes of incubation at 4 °C on an overhead rotation system, the resin was collected by centrifugation at 1000 g for 5 minutes at 4 °C. The extraction was then washed four times with a wash buffer (50 mM HEPES-KOH (pH 7.2), 5% glycerol and 300 mM NaCl, 1x protease inhibitor cocktail). The proteins of interest were then eluted using a maltose elution buffer (50 mM HEPES-KOH (pH7.2), 5% glycerol, 100 mM NaCl and 200 mM maltose, 2 mM DTT). To exchange the buffer in which the proteins were eluted, the samples were run through 50 kDa exclusion centrifuge filters (Amicon, Ultra Zentrifugenfilter, 50 kDa MWCO) and diluted with the elution buffer lacking maltose. MBP-AvrPm4^96224^ (expected size 80 kDa) migrated as a band at around 110kDa, with degradation products ranging from 110 to 70kDa, while MBP-avrPm4^4AB-2^ (expected size 67 kDa) appeared as a strong band at 85kDa, and a minor band at 70kDa (**Supp. Fig. S5B**). Removal of the MBP tag from the AvrPm4 constructs resulted in insoluble protein. Therefore, we used tagged AvrPm4 proteins for the experiments (**Supp. Fig. S5B**).

Expression and purification of the N-terminally His-MBP-tagged Exon1–5 construct were carried out following the protocol described by (Caro *et al*., 2020), with some modifications. For this construct, the lysis buffer consisted of 50 mM Tris-HCl (pH 7.5), 200 mM NaCl, 10% glycerol, 5 mM imidazole, 1 mM PMSF, and 1× protease inhibitor cocktail (Roche, Basel, Switzerland). For the purification procedure, HisPur™ Cobalt Resin (Thermo Fisher Scientific, Waltham, MA, USA) was used. The wash buffer contained 50 mM Tris-HCl (pH 7.5), 300 mM NaCl, 10 mM imidazole, and 1 mM PMSF. Proteins were eluted from the resin at 4 °C using elution buffer composed of 50 mM Tris-HCl (pH 7.5), 200 mM NaCl, 10% glycerol, and 250 mM imidazole. Protein purity and concentration were assessed by SDS-PAGE, using bovine serum albumin (BSA) as a standard. The gels were stained with Coomassie Brilliant Blue R-250 and protein concentrations were estimated based band intensity analysis using Image J.

For the *in vitro* kinase assay, 1 µg of purified protein was incubated in a 20 µL reaction containing 1 µCi γ32P-ATP, 10 mM MgCl2, 10 mM MnCl2, 50 mM HEPES-KOH (pH 7.2), 5% glycerol, 100 mM NaCl 2 mM ATP for 60 minutes at 37 °C. Proteins were separated by SDS-PAGE and transferred onto a PVDF membrane, which was then exposed overnight to a phosphor screen before imaging using the Amersham Typhoon system (GE Healthcare Life Sciences, Chicago, IL, USA).

### Protein detection

Immunoblotting was initiated by blocking the membrane with 5% (w/v) non-fat dry milk in TBS-T. MBP-tagged proteins were detected using anti-MBP antibody (New England Biolabs, Ipswich, MA, USA; IgG2a, E8032S) at a 1:3,000 dilution. For LUC-tagged proteins, a polyclonal anti-luciferase antibody (Sigma-Aldrich, St. Louis, MO, USA; L0159) was used at 1:3,000 dilution, followed by a secondary anti-rabbit HRP-conjugated antibody (LabForce, Nunningen, Switzerland; sc-2357) also at 1:3,000 dilution. Detection of mTurquoise and Venus-tagged proteins was carried out using anti-GFP antibody (clone B-2; Santa Cruz Biotechnology, Dallas, TX, USA; sc-9996) at a 1:5,000 dilution. mCherry-tagged proteins were detected using anti-RFP antibody (clone 6G6; Chromotek, Planegg, Germany) at a 1:5,000 dilution. In both cases, the corresponding secondary antibody was an anti-mouse HRP-conjugated antibody (Promega, Madison, WI, USA; W402B) at 1:10,000 dilution. Peroxidase chemiluminescence was detected using a Fusion FX Imaging System (Vilber Lourmat, Eberhardzell, Germany) after application of Western Bright ECL HRP substrate (Advansta, Menlo Park, CA, USA). The SuperSignal™ West Femto substrate (Thermo Scientific, Waltham, MA, USA) was used when maximum sensitivity was required.

### HR measurement in *N. benthamiana*

To assess HR responses upon co-expression of effector and R genes, *Agrobacterium* cultures were prepared as described above and mixed in a 4:1 ratio (effector:R) prior to infiltration. Four to five days post infiltration, imaging and quantification of HR responses was achieved using a Fusion FX Imaging System (Vilber Lourmat, Eberhardzell, Germany) and the ImageJ software as described by (Bourras *et al*., 2019).

### Total protein extraction from *N. benthamiana*

*N. benthamiana* tissue for total protein extraction was harvested 3 days after *Agrobacterium* infiltration by cutting ten leaf disks (6mm diameter) from 3 independent plants and immediately frozen in liquid nitrogen. The samples were collected in a 2 mL tube with 3 glass beads and next ground using a tissue homogeniser. 1.5x Lämmli buffer was added and the samples incubated at 95 °C for 10 min. After 1 min centrifugation at 13,000 rpm the supernatant was used to run an SDS-PAGE.

### Structural modelling and other bioinformatic analyses

Domain prediction of AvrPm4^96224^ was performed with NCBI CD search (available through https://www.ncbi.nlm.nih.gov/Structure/cdd/wrpsb.cgi). Modelling of AvrPm4^96224^ has been performed using Alphafold3, available through https://alphafoldserver.com/. The dotplot of AvrPm4^96224^ has been done and visualised using an in-house software by T. Wicker. Protein alignments have been performed with ClustalOmega and visualised using CLC Main Workbench. The nuclear localisation signal was predicted using NLStradamus (http://www.moseslab.csb.utoronto.ca/NLStradamus/). The prediction of the DNA binding probability was performed using DP-bind (https://lcg.rit.albany.edu/dp-bind/) with standard parameters; the raw data are in **Supp. Table S6**. Bins of 10 amino acid residues containing their average value (between 0 = non-binding and 1 = binding) were created and used for producing the density plot depicted in **Supp. Fig. 3D**.

### Statistical analyses

All statistical analyses were performed in R/Rstudio (versions 4.3.1 and 2023.06.1, respectively). To evaluate differences between treatments in all protoplast and *N. benthamiana* cell death assays, analysis of variance (ANOVA) followed by Tukey HSD tests were performed (R package multcomp). For the split-luciferase assay, an ANOVA followed by multiple comparison of means was performed. To assign letters, the commands *glht* and *cld* from the R package *multcomp* have been used. Package *ggplot2* was used for producing graphs. Raw data are in **Supp. Table S7**.

### GWAS analysis

For the GWAS analysis, Illumina resequencing data was mapped against the *Bgt* reference genome (Bgt_genome_v3_16) following the procedure described in (Kunz *et al*., 2023). Single nucleotide polymorphisms (SNPs) were identified using the FreeBayes tool with the command freebayes -p 1 (v1.3.6, (Garrison & Marth, 2012)). The identified polymorphic sites were then filtered using VCFtools (v0.1.16, (Danecek *et al*., 2011)) with the following specifications: --max-alleles 2 --min-alleles --maf 0.05 –max-missing 0.95, --minDP 8, and transformed to hapmap format using a custom perl script. Subsequently, GWAS analysis was conducted with virulence phenotyping data of 78 *Bgt* isolates on NILs Khapli/8*CC (*Pm4a*) and W804/8*Fed (*Pm4b*) (**Supp. Table S3**) using the GAPIT software (v3, (Wang & Zhang, 2021)) with the following specifications; PCA.total = 3, model =c(“MLM”).

To estimate the interval underlying a significant association in the GWAS analysis, the pairwise R² values between markers on Chr-04 and Chr-08 with less than 1M bp distance were determined using the VCFtools commands --ld-window-bp 1000000 --max-alleles 2 --min-alleles 2 --geno-r2 (v0.1.16, (Danecek *et al*., 2011)). Finally, SNPs with a R^2^ value greater than 0.6 upon pairwise comparison with most significantly associated SNPs of the GWAS analysis were used to define the genomic candidate interval.

### QTL mapping

QTL mapping was performed based on the previously described genetic map of the cross between THUN-12 (*B.g. triticale*) X CHE_96224 (*Bgt*) ((Müller *et al*., 2019; Hewitt *et al*., 2020)) which is available from https://github.com/MarionCMueller. The QTL analysis was performed using the r/qtl package in R (v1.70, (Broman *et al*., 2003)). First, the genetic map was processed using the command jittermap() and calc.genoprob(step=2,error.prob = 0.001), followed by QTL analysis using the command scanone(model = “np”). The significance threshold at alpha=0.05 was determined via calculating 1000 permutation with scanone(). The 1.5LOD interval was determined using lodint().

### Candidate identification and expression analysis of *SvrPm4* candidates

To identify *SvrPm4* candidate genes, candidate interval on Chr-08 was first inspected for spurious annotations. Two gene models were detected, *Bgt-51527* and *Bgt-20620-4*, which either lacked a start or stop codon. Consequently, these genes were excluded from subsequent analysis. Next, sequence variants of the three candidate effectors located in the interval (*Bgt-51526, BgtAcSP-30824* and *BgtAcSP-31023*) were detected in a set of reference isolates (CHE_96224, CHE_94202, GBR_JIW2, and ISR_7) based on resequencing data (**Supp. Table S3**).

To accurately determine the expression levels of the three candidate effector genes in the isolates CHE_96224, CHE_94202, GBR_JIW2, and ISR_7, RNA sequencing data from these isolates on the susceptible wheat cultivar ‘Chinese Spring’ was used at two days post inoculation (Praz *et al*., 2018; Kunz *et al*., 2023). To quantify transcript abundance, the annotation of CHE_96224 was used (available at https://zenodo.org/records/7018501). To avoid quantification biases, CDS sequences of all gene variants present in the four reference isolates to the CHE_96224 CDS annotation file were added. Finally, the new Fasta file was processed using the salmon index command from the salmon package (v1.4.0, (Patro *et al*., 2017)), with the chromosomes of the Bgt_genome_v3_16 assemblies as decoys. Transcripts were quantified transcripts using the salmon quant -l A command. Subsequently, expression data across all variants for each isolate was merged in R and calculated RPKM values using the rpkm() function from the edgeR (v4.0.16, (Robinson *et al*., 2010)).

## Acknowledgements

We thank Matthias Heuberger for bioinformatics support, Gerhard Herren for technical support, Karl Huwiler and Urs Somalvico for greenhouse support.

## Author contributions

Z.B., A.H., M.D.P.C., L.K., J.S.-M., and B.K. conceived the experiments, interpreted results, and wrote the manuscript. Z.B. and U.S. identified and analysed the *Bgt* mutants. A.H., M.D.P.C. and Z.B. performed the protoplast cell death assays. Z.B. performed the protein-protein interaction assay.

M.D.P.C. performed confocal microscopic analysis. A.H., M.D.P.C. and Z.B. performed protein extraction and purification. M.N., A.H., Z.B., M.D.P.C, V.W., and J.S.-M. performed the kinase assays. L.K., M.C.M. and S.S. performed the GWAS analysis and QTL mapping and interpreted results. L.K. performed HR measurements in *N. benthamiana*. S.R. and J.I. were involved in cloning and different experimental procedures. K.B. and C.Z. provided support in establishing the in vitro kinase assay. M.O., M.F. and P.D. supported the establishment of the protoplast cell death assay. T.W. provided support in bioinformatic analyses. All authors have read and agreed to the published version of the article.

## Data availability

Raw FASTA sequences of all *Bgt* mutants generated in this study are available through NCBI (BioProject ID: PRJNA1016363).

## References

Anh V, Inoue Y, Asuke S, Vy T, Anh N, Wang S, Chuma I, Tosa Y. 2018. Rmg8 and Rmg7, wheat genes for resistance to the wheat blast fungus, recognize the same avirulence gene AVR-Rmg8. MOLECULAR PLANT PATHOLOGY 19(5): 1252–1256.

Arndell T, Chen J, Sperschneider J, Upadhyaya N, Blundell C, Niesner N, Outram M, Wang A, Swain S, Luo M, et al. 2024. Pooled effector library screening in protoplasts rapidly identifies novel Avr genes. NATURE PLANTS 10(4).

Arora S, Steed A, Goddard R, Gaurav K, O’Hara T, Schoen A, Rawat N, Elkot A, Korolev A, Chinoy C, et al. 2023. A wheat kinase and immune receptor form host-specificity barriers against the blast fungus. NATURE PLANTS 9(3): 385–392.

Arora S, Steuernagel B, Gaurav K, Chandramohan S, Long YM, Matny O, Johnson R, Enk J, Periyannan S, Singh N, et al. 2019. Resistance gene cloning from a wild crop relative by sequence capture and association genetics. Nature Biotechnology 37(2): 139–143.

Asuke S, Morita K, Shimizu M, Abe F, Terauchi R, Nago C, Takahashi Y, Shibata M, Yoshioka M, Iwakawa M, et al. 2024. Evolution of wheat blast resistance gene Rmg8 accompanied by differentiation of variants recognizing the powdery mildew fungus. NATURE PLANTS 10(6).

Bernasconi Z, Stirnemann U, Heuberger M, Sotiropoulos A, Graf J, Wicker T, Keller B, Sanchez-Martin J. 2024. Mutagenesis of Wheat Powdery Mildew Reveals a Single Gene Controlling Both NLR and Tandem Kinase-Mediated Immunity. MOLECULAR PLANT-MICROBE INTERACTIONS 37(3): 264–276.

Bilstein-Schloemer M, Muller MC, Saur IML. 2025. Technical Advances Drive the Molecular Understanding of Effectors from Wheat and Barley Powdery Mildew Fungi. Molecular plant-microbe interactions : MPMI 38(2): 213–225.

Bolger AM, Lohse M, Usadel B. 2014. Trimmomatic: a flexible trimmer for Illumina sequence data. Bioinformatics 30(15): 2114–2120.

Bourras S, Kunz L, Xue MF, Praz CR, Muller MC, Kalin C, Schlafli M, Ackermann P, Fluckiger S, Parlange F, et al. 2019. The AvrPm3-Pm3 effector-NLR interactions control both race-specific resistance and host-specificity of cereal mildews on wheat. Nature Communications 10(1): 2292.

Bourras S, McNally KE, Ben-David R, Parlange F, Roffler S, Praz CR, Oberhaensli S, Menardo F, Stirnweis D, Frenkel Z, et al. 2015. Multiple avirulence loci and allele-specific effector recognition control the Pm3 race-specific resistance of wheat to powdery mildew. Plant Cell 27(10): 2991–3012.

Bourras S, McNally KE, Mueller MC, Wicker T, Keller B. 2016. Avirulence genes in cereal powdery mildews: the gene-for-gene hypothesis 2.0. Frontiers in Plant Science 7: 241.

Brabham H, De La Cruz D, Were V, Shimizu M, Saitoh H, Hernández-Pinzón I, Green P, Lorang J, Fujisaki K, Sato K, et al. 2024. Barley MLA3 recognizes the host-specificity effector Pwl2 from Magnaporthe oryzae. PLANT CELL 36(2): 447–470.

Broman KW, Wu H, Sen S, Churchill GA. 2003. R/qtl: QTL mapping in experimental crosses. Bioinformatics 19(7): 889–890.

Brunner S, Hurni S, Streckeisen P, Mayr G, Albrecht M, Yahiaoui N, Keller B. 2010. Intragenic allele pyramiding combines different specificities of wheat Pm3 resistance alleles. Plant Journal 64(3): 433–445.

Burgess A, Buck M, Krauer K, Sculley T. 2006. Nuclear localization of the Epstein-Barr virus EBNA3B protein. JOURNAL OF GENERAL VIROLOGY 87: 789–793.

Caro M, Holton N, Conti G, Venturuzzi A, Martínez-Zamora M, Zipfel C, Asurmendi S, Díaz-Ricci J. 2020. The fungal subtilase AsES elicits a PTI-like defence response in Arabidopsis thaliana plants independently of its enzymatic activity. MOLECULAR PLANT PATHOLOGY 21(2): 147–159.

Chen JP, Upadhyaya NM, Ortiz D, Sperschneider J, Li F, Bouton C, Breen S, Dong CM, Xu B, Zhang XX, et al. 2017. Loss of *AvrSr50* by somatic exchange in stem rust leads to virulence for *Sr50* resistance in wheat. Science 358(6370): 1607–1610.

Chen R, Chen J, Powell O, Outram M, Arndell T, Gajendiran K, Wang Y, Lubega J, Xu Y, Ayliffe M, et al. 2025. A wheat tandem kinase activates an NLR to trigger immunity. SCIENCE 387(6741): 1402–1408.

Chen R, Gajendiran K, Wulff B. 2024. R we there yet? Advances in cloning resistance genes for engineering immunity in crop plants. CURRENT OPINION IN PLANT BIOLOGY 77.

Danecek P, Auton A, Abecasis G, Albers CA, Banks E, DePristo MA, Handsaker RE, Lunter G, Marth GT, Sherry ST, et al. 2011. The variant call format and VCFtools. Bioinformatics 27(15): 2156–2158.

Dodds PN, Lawrence GJ, Catanzariti AM, Ayliffe MA, Ellis JG. 2004. The Melampsora lini AvrL567 avirulence genes are expressed in haustoria and their products are recognized inside plant cells. Plant Cell 16(3): 755–768.

Dodds PN, Rathjen JP. 2010. Plant immunity: towards an integrated view of plant-pathogen interactions. Nature Reviews Genetics 11(8): 539–548.

Fu D, Uauy C, Distelfeld A, Blechl A, Epstein L, Chen X, Sela H, Fahima T, Dubcovsky J. 2009. A Kinase-START Gene Confers Temperature-Dependent Resistance to Wheat Stripe Rust. SCIENCE 323(5919): 1357–1360.

Garrison E, Marth G. 2012. Haplotype-based variant detection from short-read sequencing. arXiv: 1207.3907v1202.

Gehl C, Kaufholdt D, Hamisch D, Bikker R, Kudla J, Mendel R, Hänsch R. 2011. Quantitative analysis of dynamic protein-protein interactions in planta by a floated-leaf luciferase complementation imaging (FLuCI) assay using binary Gateway vectors. PLANT JOURNAL 67(3): 542–553.

Ghanbarnia K, Ma L, Larkan N, Haddadi P, Fernando W, Borhan M. 2018. Leptosphaeria maculans AvrLm9: a new player in the game of hide and seek with AvrLm4-7. MOLECULAR PLANT PATHOLOGY 19(7): 1754–1764.

Hewitt T, Müller MC, Molnár I, Mascher M, Holušová K, Šimková H, Kunz L, Zhang J, Li J, Bhatt D, et al. 2020. A highly differentiated region of wheat chromosome 7AL encodes a *Pm1*a immune receptor that recognises its corresponding *AvrPm1a* effector from *Blumeria graminis*. New Phytologist 229(5): 2812–2826.

Himmelbach A, Zierold U, Hensel G, al. e. 2007. A set of modular binary vectors for transformation of cereals. Plant Physiol. 145(4): 1192–1200.

Hurni S, Brunner S, Buchmann G, Herren G, Jordan T, Krukowski P, Wicker T, Yahiaoui N, Mago R, Keller B. 2013. Rye *Pm8* and wheat *Pm3* are orthologous genes and show evolutionary conservation of resistance function against powdery mildew. Plant Journal 76(6): 957–969.

Inoue Y, Vy TTP, Tani D, Tosa Y. 2021. Suppression of wheat blast resistance by an effector ofPyricularia oryzaeis counteracted by a host specificity resistance gene in wheat. New Phytologist 229(1): 488–500.

Jay F, Brioudes F, Voinnet O. 2023. A contemporary reassessment of the enhanced transient expression system based on the tombusviral silencing suppressor protein P19. PLANT JOURNAL 113(1): 186–204.

Kloppe T, Whetten RB, Kim S-B, Powell OR, Luck S, Douchkov D, Whetten RW, Hulse-Kemp AM, Balint-Kurti P, Cowger C. 2023. Two pathogen loci determine Blumeria graminis f. sp. tritici virulence to wheat resistance gene Pm1a. New Phytologist 238(4): 1546–1561.

Koller T, Brunner S, Herren G, Sanchez-Martin J, Hurni S, Keller B. 2019. Field grown transgenic Pm3e wheat lines show powdery mildew resistance and no fitness costs associated with high transgene expression. TRANSGENIC RESEARCH 28(1): 9–20.

Krauer K, Belzer D, Liaskou D, Buck M, Cross S, Honjo T, Sculley T. 1998. Regulation of interleukin-1β transcription by Epstein-Barr virus involves a number of latent proteins via their interaction with RBP. VIROLOGY 252(2): 418–430.

Kunz L, Sotiropoulos AG, Graf J, Razavi M, Keller B, Müller MC. 2023. The broad use of the Pm8 resistance gene in wheat resulted in hypermutation of the AvrPm8 gene in the powdery mildew pathogen. BMC Biology 21(1): 29.

Li H, Durbin R. 2009. Fast and accurate short read alignment with Burrows-Wheeler transform. Bioinformatics 25(14): 1754–1760.

Li H, Handsaker B, Wysoker A, Fennell T, Ruan J, Homer N, Marth G, Abecasis G, Durbin R, Genome Project Data P. 2009. The Sequence Alignment/Map format and SAMtools. Bioinformatics 25(16): 2078–2079.

Liu Y, Hou S, Chen S. 2024. Kinase fusion proteins: intracellular R-proteins in plant immunity. TRENDS IN PLANT SCIENCE 29(3): 278–282.

Lu P, Guo L, Wang Z, Li B, Li J, Li Y, Qiu D, Shi W, Yang L, Wang N, et al. 2020. A rare gain of function mutation in a wheat tandem kinase confers resistance to powdery mildew. Nature Communications 11(1): 680.

Lu P, Zhang G, Li J, Gong Z, Wang G, Dong L, Zhang H, Guo G, Su M, Wang K, et al. 2025. A wheat tandem kinase and NLR pair confers resistance to multiple fungal pathogens. SCIENCE 387(6741): 1418–1424.

Malik S, Roeder R. 2010. The metazoan Mediator co-activator complex as an integrative hub for transcriptional regulation. NATURE REVIEWS GENETICS 11(11): 761–772.

Müller MC, Praz CR, Sotiropoulos AG, Menardo F, Kunz L, Schudel S, Oberhänsli S, Poretti M, Wehrli A, Bourras S, et al. 2019. A chromosome-scale genome assembly reveals a highly dynamic effector repertoire of wheat powdery mildew. New Phytologist 221(4): 2176–2189.

Ngou BPM, Ding P, Jones JDG. 2022. Thirty years of resistance: Zig-zag through the plant immune system. Plant Cell 34(5): 1447–1478.

Nirmala J, Drader T, Lawrence P, Yin C, Hulbert S, Steber C, Steffenson B, Szabo L, von Wettstein D, Kleinhofs A. 2011. Concerted action of two avirulent spore effectors activates Reaction to Puccinia graminis 1 (Rpg1)-mediated cereal stem rust resistance. PROCEEDINGS OF THE NATIONAL ACADEMY OF SCIENCES OF THE UNITED STATES OF AMERICA 108(35): 14676–14681.

O’Hara T, Steed A, Goddard R, Gaurav K, Arora S, Quiroz-Chávez J, Ramirez-González R, Badgami R, Gilbert D, Sánchez-Martín J, et al. 2024. The wheat powdery mildew resistance gene Pm4 also confers resistance to wheat blast. NATURE PLANTS 10(6).

Parlange F, Oberhaensli S, Breen J, Platzer M, Taudien S, Simkova H, Wicker T, Dolezel J, Keller B. 2011. A major invasion of transposable elements accounts for the large size of the Blumeria graminis f.sp tritici genome. Functional & Integrative Genomics 11(4): 671–677.

Patro R, Duggal G, Love MI, Irizarry RA, Kingsford C. 2017. Salmon provides fast and bias-aware quantification of transcript expression. Nature Methods 14(4): 417–419.

Petersen TN, Brunak S, von Heijne G, Nielsen H. 2011. SignalP 4.0: discriminating signal peptides from transmembrane regions. Nature Methods 8(10): 785–786.

Petit-Houdenot Y, Fudal I. 2017. Complex Interactions between Fungal Avirulence Genes and Their Corresponding Plant Resistance Genes and Consequences for Disease Resistance Management. Frontiers in Plant Science 8.

Plissonneau C, Daverdin G, Ollivier B, Blaise F, Degrave A, Fudal I, Rouxel T, Balesdent M. 2016. A game of hide and seek between avirulence genes AvrLm4-7 and AvrLm3 in Leptosphaeria maculans. NEW PHYTOLOGIST 209(4): 1613–1624.

Praz CR, Menardo F, Robinson MD, Müller MC, Wicker T, Bourras S, Keller B. 2018. Non-parent of origin expression of numerous effector genes indicates a role of gene regulation in host adaption of the hybrid triticale powdery mildew pathogen. Frontiers in Plant Science 9: 49.

Reveguk T, Fatiukha A, Potapenko E, Reveguk I, Sela H, Klymiuk V, Li Y, Pozniak C, Wicker T, Coaker G, et al. 2025. Tandem kinase proteins across the plant kingdom. NATURE GENETICS 57(1).

Robinson MD, McCarthy DJ, Smyth GK. 2010. edgeR: a Bioconductor package for differential expression analysis of digital gene expression data. Bioinformatics 26(1): 139–140.

Saur IML, Bauer S, Kracher B, Lu XL, Franzeskakis L, Muller MC, Sabelleck B, Kummel F, Panstruga R, Maekawa T, et al. 2019. Multiple pairs of allelic MLA immune receptor-powdery mildew AVR(A) effectors argue for a direct recognition mechanism. Elife 8: e44471.

Savary S, Willocquet L, Pethybridge SJ, Esker P, McRoberts N, Nelson A. 2019. The global burden of pathogens and pests on major food crops. Nature Ecology & Evolution 3(3): 430–439.

Singh SP, Hurni S, Ruinelli M, Brunner S, Sánchez-Martín J, Krukowski P, Peditto D, Buchmann G, Zbinden H, Keller B. 2018. Evolutionary divergence of the rye *Pm17* and *Pm8* resistance genes reveals ancient diversity. Plant Molecular Biology 98(3): 249–260.

Sotiropoulos AG, Arango-Isaza E, Ban T, Barbieri C, Bourras S, Cowger C, Czembor PC, Ben-David R, Dinoor A, Ellwood SR, et al. 2022. Global genomic analyses of wheat powdery mildew reveal association of pathogen spread with historical human migration and trade. Nature communications 13(1): 4315–4315.

Sung Y, Li Y, Bernasconi Z, Baik S, Asuke S, Keller B, Fahima T, Coaker G. 2025. Wheat tandem kinase RWT4 directly binds a fungal effector to activate defense. NATURE GENETICS.

Sánchez-Martín J, Keller B. 2021. NLR immune receptors and diverse types of non-NLR proteins control race-specific resistance in Triticeae. Current Opinion in Plant Biology 62: 102053.

Sánchez-Martín J, Steuernagel B, Ghosh S, Herren G, Hurni S, Adamski N, Vrana J, Kubalakova M, Krattinger SG, Wicker T, et al. 2016. Rapid gene isolation in barley and wheat by mutant chromosome sequencing. Genome Biology 17: 221.

Sánchez-Martín J, Widrig V, Herren G, Wicker T, Zbinden H, Gronnier J, Spörri L, Praz CR, Heuberger M, Kolodziej MC, et al. 2021. Wheat Pm4 resistance to powdery mildew is controlled by alternative splice variants encoding chimeric proteins. Nature Plants 4: 327–341.

Walker I, Hsieh P, Riggs P. 2010. Mutations in maltose-binding protein that alter affinity and solubility properties. APPLIED MICROBIOLOGY AND BIOTECHNOLOGY 88(1): 187–197.

Wang J, Zhang Z. 2021. GAPIT Version 3: Boosting Power and Accuracy for Genomic Association and Prediction. Genomics Proteomics & Bioinformatics 19(4): 629–640.

Wu C, Derevnina L. 2023. The battle within: How pathogen effectors suppress NLR-mediated immunity. CURRENT OPINION IN PLANT BIOLOGY 74.

Xie J, Luo Q, Wang L, Qiu D, Zhao C, Hu J, Zhang J, Zhao X, Chen Z, Wang Y, et al. 2025. A biGWAS strategy reveals the genetic architecture of the interaction between wheat and *Blumeria graminis* f.sp. *tritici*. bioRxiv: doi:10.1101/2025.1104.1109.647224.

