## Supplementary Figures for "Virulence on Pm4 kinase-based resistance is determined by two divergent wheat powdery mildew effectors"

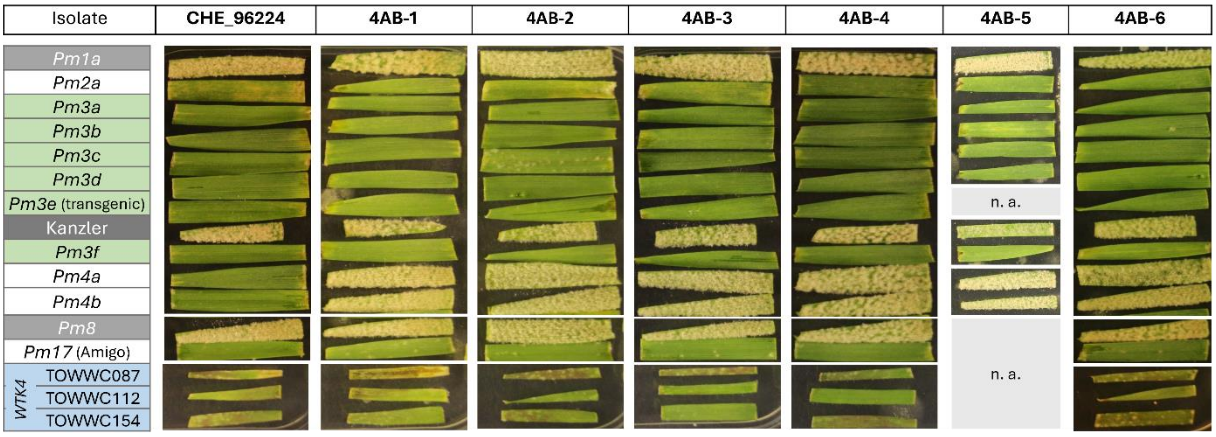


**Supp. Fig. S1. *AvrPm4* mutants are virulent on Pm4-containing lines.** The *AvrPm4* mutants exhibit specific gain of virulence on *Pm4a* and *Pm4b* near isogenic lines (NILs) but no virulence on various other *R* genes for which their parental isolate CHE_96224 is avirulent. Mutant 4AB-5 died shortly after sequencing and it was not possible to perform phenotyping on all *Pm* lines. Cultivars susceptible to CHE_96224 (the *Pm1a* NIL, the *Pm8* NIL and Kanzler) are highlighted in grey. The NILs containing *Pm3* alleles are depicted in green, and the three *Ae. tauschii* lines containing *WTK4* are in blue. The remaining *Pm* lines are shown in white.

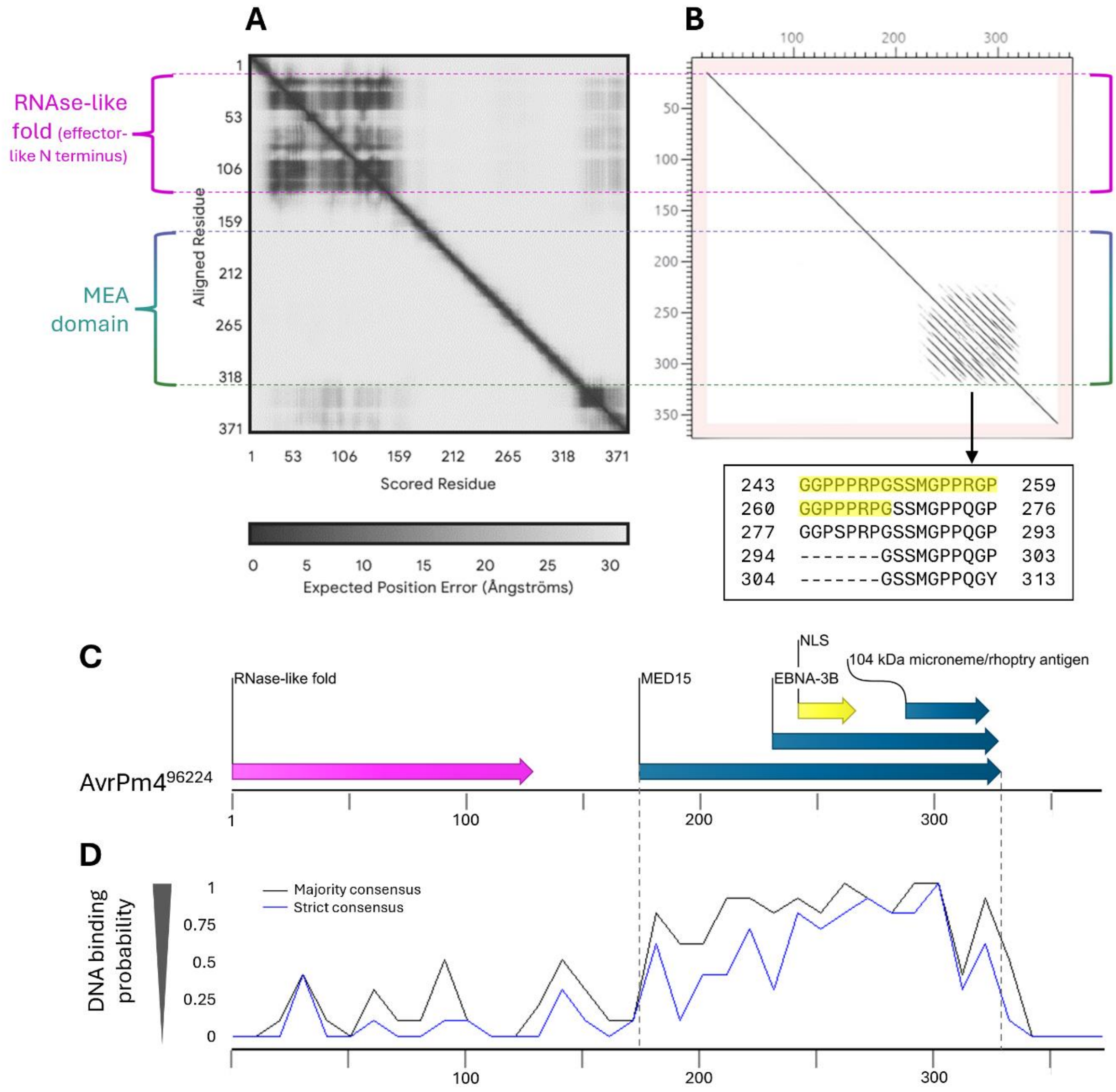


**Supp. Fig. S2. AvrPm4 has an unstructured, highly repetitive, DNA binding MEA domain**. (**A**) The predicted aligned error (PAE) of the Alphafold prediction (shown in **Fig. 1C**) highlights a high quality of the prediction (low error) in the effector-like N terminus with an RNase-like fold (in pink), and a poor prediction of the MEA domain (in blue/green). (**B**) A dot plot (visual alignment) of AvrPm4$^{96224}$ against itself at protein level reveals multiple duplicated fragments in the second half of the MEA domain. The duplicated fragments are 17 (or 10) amino acids long, glycine and proline rich, and contain a nuclear localisation signal (NLS) predicted by NLStradamus (in yellow). (**C**) The domain prediction from NCBI conserved domain search reveals three overlapping hits (in blue), with the MED15 domain being the largest (aa 175-329), containing an EBNA-3B (aa 231-328) and a 104 kDa microneme/rhoptry antigen (aa 288-324) domains, with E-values of 4.23e$^{-3}$, 2.83e$^{-3}$ and 1.07e$^{-3}$, respectively. The NLS (aa 243-267) is contained in both the EBNA-3B and the MED15 domain (in yellow). The first 129 amino acids have an RNase-like fold (in pink). (**D**) Density plot of DNA binding probability of AvrPm4 predicted by DP-bind. Binding (value = 1) and non-binding amino acid residues (value = 0) were binned in bins of 10 amino acids, and the average value of the bin is depicted in the plot. The black line indicates the majority consensus (at least 2 of the 3 DP-bind models predict the residue to bind DNA) and the blue line indicates the strict consensus (all 3 models predict the residue to be DNA binding). Raw data from DP-bind are in **Supp. Table S6**.

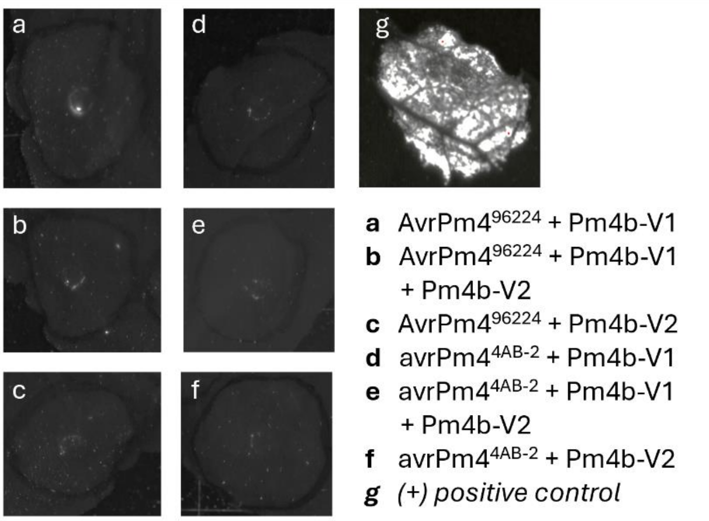


**Supp. Fig. S3. *AvrPm4$^{96224}$* does not induce cell death in co-expression with *Pm4* in *N. benthamiana***. The cell death signal was measured 6 days post infiltration. For all treatments, 10% P19 was co-infiltrated. At least 3 replicates per treatment were infiltrated.

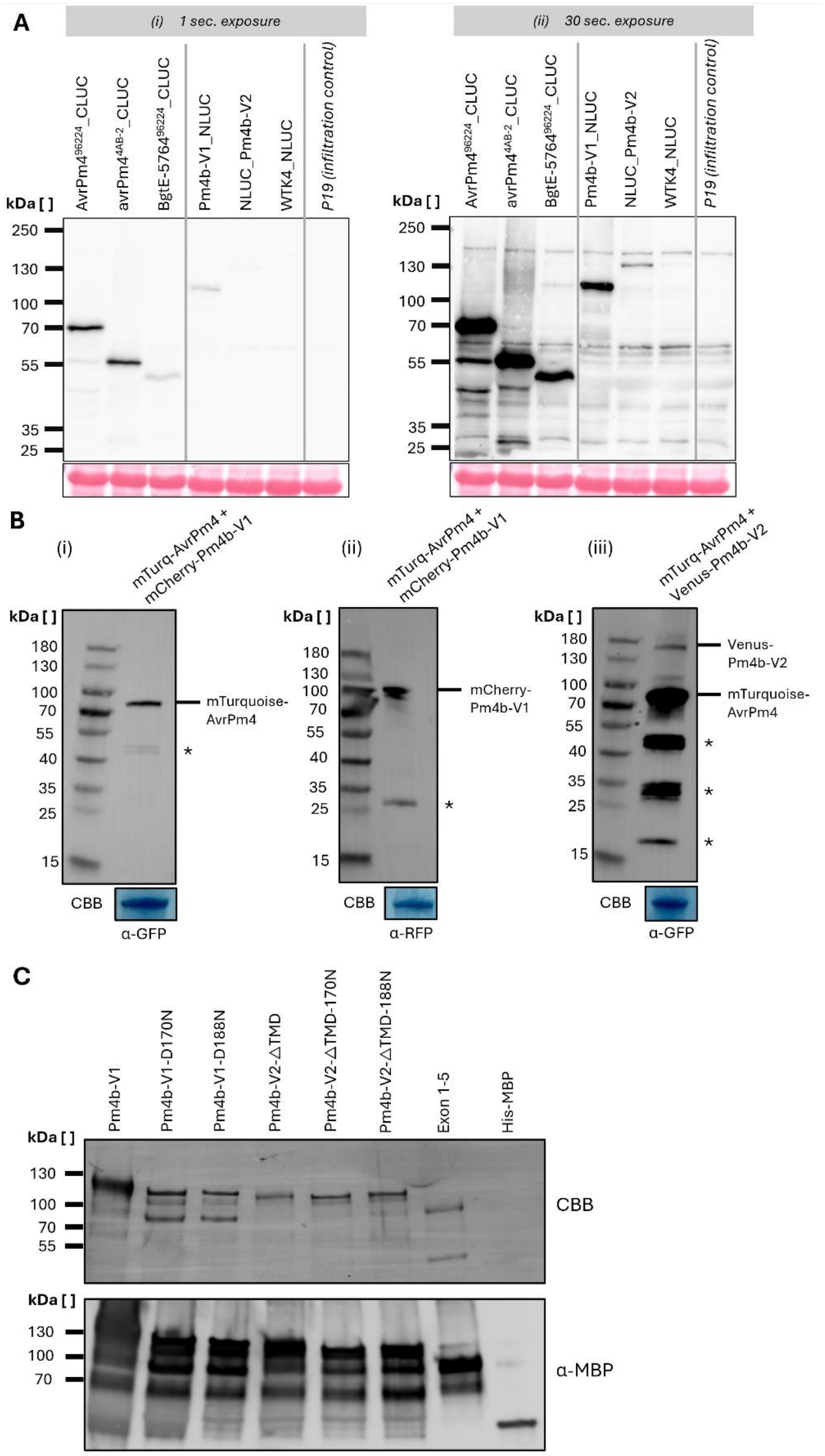


**Supp. Fig. S4. Western blots of different AvrPm4 and Pm4 constructs.** (**A**) Protein levels of the different constructs used for the split luciferase experiment (**Fig. 3A-B**). The same membrane was imaged for 1 second (i) or 30 seconds (ii) exposure time. Ponceau staining below the blots shows equal protein loading. (**B**) Detection of full-length mTurquoise-AvrPm4 (65 kDa) co-expressed with either mCherry-Pm4b_V1 (90 kDa) or Venus-Pm4b-V2 (113 kDa) in the *N. benthamiana* samples used for the co-localisation experiments. The unlabelled bands observed in the WB (indicated with an asterisk) correspond to partial degradation products of the target protein and/or free fluorophore. (**C**) Coomassie Brilliant Blue G-250 staining and western blot of purified MBP-tagged proteins used in the in-vitro kinase assay.

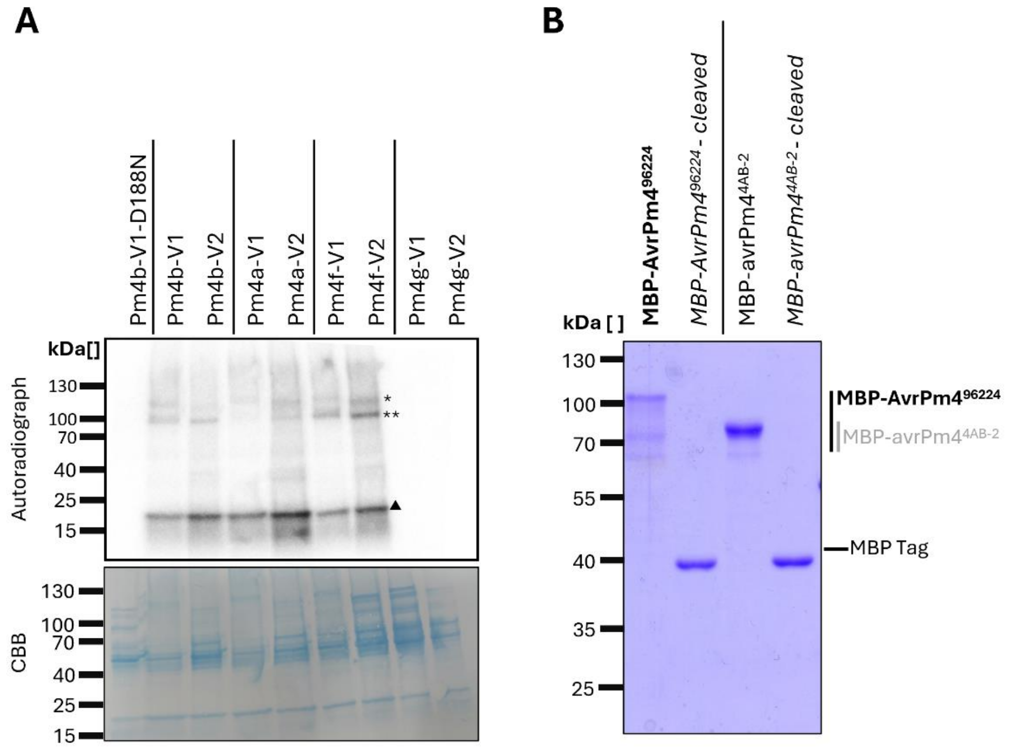


**Supp. Fig. S5. Different Pm4 variants show different kinase activity and AvrPm4 precipitates without MBP tag. (A)** *In vitro* autophosphorylation and transphosphorylation assays in the presence of [ɣ-32P] and myelin basic protein using purified Pm4-V1 and V2 of the *Pm4* alleles: P*m4a, Pm4b, Pm4f,* and *Pm4g*. All constructs shown have an MBP tag. Different allelic variants show a consistently observed truncated form of Pm4, migrating between 80 kDa and 90 kDa, which likely results from proteolytic degradation. Coomassie Brilliant Blue R-250 stained PVDF membrane is indicated as a loading control. (**B**) MBP tagged AvrPm4 was purified as described in material and methods. An aliquot of the purified sample was incubated with enterokinase to cleave off the MBP tag. The four samples were subsequently run on an SDS-PAGE and Coomassie stained. MBP-AvrPm4$^{96224}$ (predicted size 80 kDa) migrates at 110kDa, with truncated versions between 100kDa and 70kDa. MBP-avrPm4$^{4AB-2}$ (67 kDa) migrates between the 70 kDa and 85 kDa mark and shows a truncated occurring version at 70kDa. After cleavage with enterokinase, the MBP tag is visible at 42 kDa. AvrPm4 is no longer detectable on the gel, likely due to its insolubility in absence of the MBP tag, resulting in precipitation.

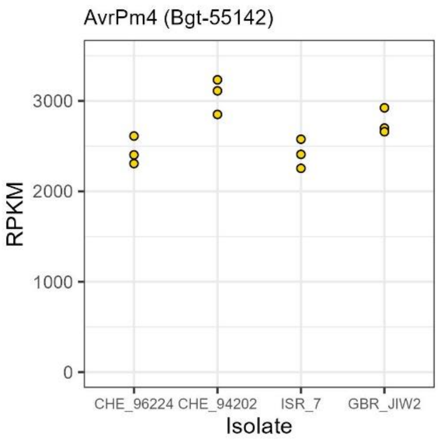


**Supp. Fig. S6. *AvrPm4* (*Bgt-55142*) expression levels in four *Bgt* reference isolates CHE_96224, CHE_94202, ISR_7 and GBR_JIW2.** Expression levels are shown as RPKM values at 48 hours post infection on the susceptible wheat cultivar 'Chinese Spring'. Values of three biological replicates are shown as yellow circles.

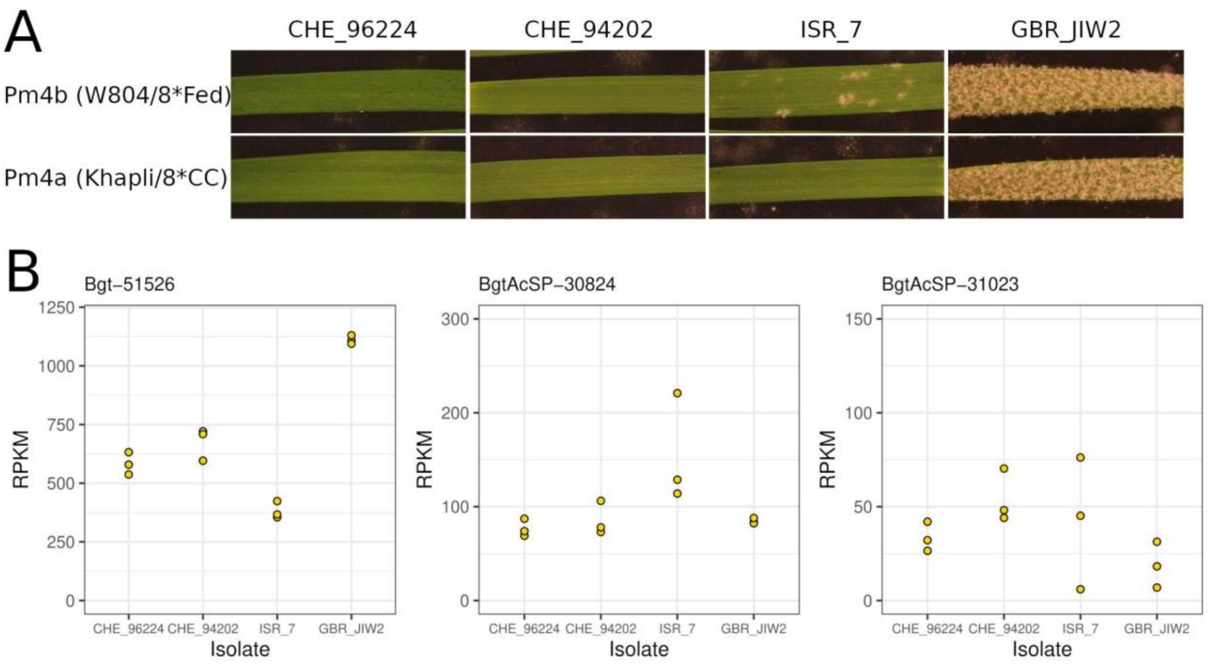


**Supp. Fig. S7. *Pm4b* and *Pm4a* phenotypes and *SvrPm4* candidate gene expression levels in four *Bgt* reference isolates.** (**A**) Virulence phenotypes of reference isolates CHE_96224, CHE_94202, ISR_7 and GBR_JIW2 on the near isogenic lines 'W804/8*Fed' (*Pm4b*) and 'Khapli/8*CC' (*Pm4a*). Pictures were taken eight days post inoculation. (**B**) Expression levels of *SvrPm4* candidate genes *Bgt-51526, BgtAcSP-30824* and *BgtAcSP-31023* in the four *Bgt* reference isolates CHE_96224, CHE_94202, ISR_7 and GBR_JIW2. Expression levels are shown as RPKM values at 48 hours post infection on the susceptible wheat cultivar 'Chinese Spring'. Values of three biological replicates are shown as yellow circles.

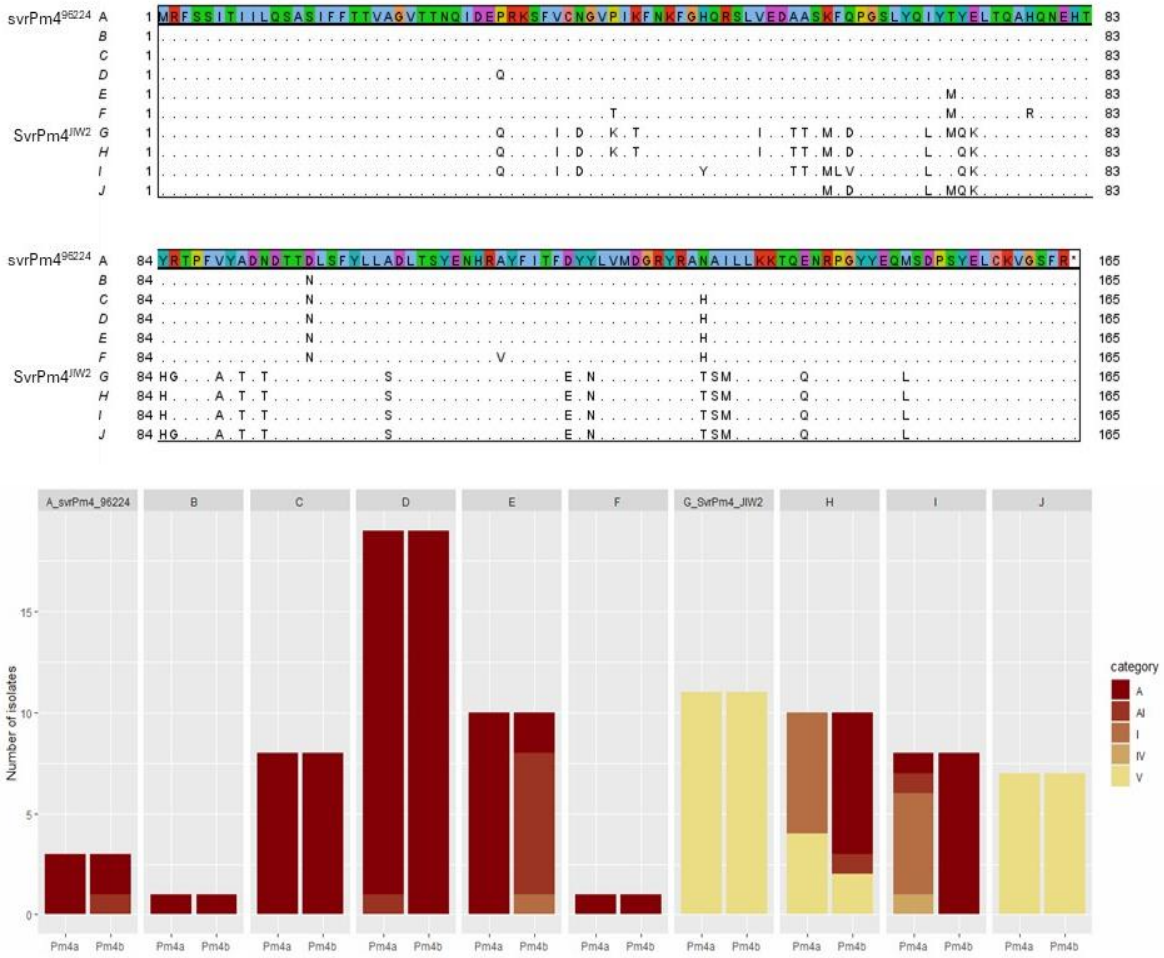


**Supp. Fig. S8. Bgt-51526 (SvrPm4) haplovariants and correlation with virulence phenotypes on *Pm4b* and *Pm4a* near-isogenic lines.** (**A**) Multiple sequence alignment of Bgt-51526 haplovariants found in the 78 tested *Bgt* isolates. Bgt-51526 protein sequence of the isolate CHE_96224 (=svrPm4$^{96224}$) is shown as a reference. Polymorphic residues in haplovariants B-J are highlighted. (**B**) Phenotypic spectrum of isolates with the indicated haplovariants on *Pm4a* and *Pm4b* NILs. Phenotypes were assessed 8 days post inoculation on at least three biological replicates and categorized based on observed mildew leaf coverage (avirulent (A) = 0%; avirulent-intermediate (AI) = (~25%); intermediate (I) = ~50%; intermediate-virulent (IV) = ~75%; virulent (V) = 100%).
